# *E. coli* extracellular matrix: a tunable composite with hierarchical structure

**DOI:** 10.64898/2026.02.22.707275

**Authors:** Macarena Siri, Agustín Mangiarotti, Anne Seewald, Nikolai Rosenthal, Shahrouz Amini, Emeline Raguin, Peter Fratzl, Cécile M. Bidan

## Abstract

*Escherichia coli* (*E. coli*) biofilms consist of bacteria, an extracellular matrix (ECM) mainly made of curli amyloid fibers, phosphoethanolamine-modified cellulose (pEtN-cellulose), and water. While curli amyloid fibers contribute to biofilm rigidity, pEtN-cellulose contributes to their cohesion. This work explores the interplay between these fibers, and how their interaction influence biofilm structure and mechanical properties. We performed a multiscale analysis on *E. coli* biofilms grown using strains producing curli and pEtN-cellulose, and only curli and only pEtN-cellulose in co-seeded ratios.

Micro-indentation experiments, confocal microscopy, and cryo-FIBSEM 3D imaging revealed a composite-like behavior of the biofilm, where its mechanical properties depend on ECM composition and organization. Spectroscopic analysis of the extracted fibers showed that their biophysical properties are influenced by their pEtN-cellulose to curli ratio and assembly.

We propose that pEtN-cellulose swelling is contrained by its interactions with rigid curli fibers. The reference *E. coli* strain maximizes this effect by assembling a curli/pEtN-cellulose hybrid material at the sub-micron scale, where its composition, interactions, and architecture can explain biofilm emergent properties. This knowledge on microbial ECM assembly opens new avenues for engineering living materials, especially for the use of bacterial biofilms as a source of bio-sourced materials.

## Introduction

Many uni- or multicellular living systems in nature are efficient biosynthetic machines that are versatile and adaptable to environmental conditions. They produce materials with interesting characteristics such as environmental responsiveness, self-assembly and self-regeneration, among others.^1,2^ These features inspire the development of engineered living materials (ELMs) performing biological functions like photosynthesis and biodegradability, in addition to their physical properties. Synthetic biologists alter the living organisms present in ELMs to program properties according to society’s need. Among these living organisms, bacteria are of great interest for their potential to be genetically engineered but also because they self-produce and self-organize a three-dimensional extracellular matrix (ECM) made of biopolymers. The resulting microbial biofilm ensures bacteria survival and is known for its stability and resilience against severe conditions. Biofilms, the interactions of their components, as well as their emergent properties are also of great interest for ELMs.^3,4^

Like many biological materials, biofilms have a hierarchical organization. They are composed of bacteria, polymeric carbohydrates, proteins (usually in the form of amyloid fibers), extracellular DNA and a large part of water that make them akin to living hydrogels.^6^ Important genetic and molecular determinants of biofilm formation have been elucidated, especially in biofilms grown at solid-air interfaces (e.g. on agar),^7–10^ such as water that greatly influences biofilm growth, structure and mechanics.^10,11^ Yet, a comprehensive understanding of their composition, their assembly, and of the interactions between their components remains a challenge.^12^ For example, *E. coli* bacteria produce curli amyloid and phosphoethanolamine cellulose (pEtN-cellulose)^8^ fibers as their main extracellular matrix (ECM) components.^13^ *E. coli* biofilm ECM has been described as a composite material – or bionanocomposite^14,15^ – where curli amyloid proteins provide rigidity and adhesion of the biofilm to organic surfaces, while pEtN-cellulose carbohydrate contributes to cohesion^15^ and mitigates the immunogenicity of curli in bacterial infection.^16^ The biophysical characteristics of curli amyloid fibers from *E. coli* bacteria were also shown to change with the biofilm growth conditions.^17,18^ In *E. coli* biofilms grown on nutritive agar, the ECM is mostly found in the middle and top parts of the biofilms – i.e. away from the nutrient source, and presents specific spatial distribution, arrangement and orientation depending on its composition.^13^ While the interactions between curli amyloid fibers and pEtN-cellulose have been shown to involve the pEtN entity and to be determining for *E. coli* biofilm architecture and mechanical properties,^12,13,19^ the exact nature of these interactions remains elusive in the complex biofilm context.^20^ Indeed, it is not clear at what length scale bacterial ECM interactions occur, what are the contributions of water in these interactions, nor what are their implications on biofilm properties emerging at other scales. Is biofilm ECM just a composite hydrogel or is it a hybrid-material, where interactions of the two main components occur at the sub-micron scale, potentially mediated by water?

In this study, we performed a multiscale exploration of the interactions between curli amyloid fibers and pEtN-cellulose, and their influence on the emergent mechanical and architectural properties of the resulting biofilms. For this, we co-seeded *E. coli* K-12 bacteria strains producing only curli (W3110) and only pEtN-cellulose (AP329) to obtain macrocolony biofilms on agar plates. We considered biofilms grown from the bacteria strain producing both curli and pEtN-cellulose (AR3110) as a reference. We then performed a multimodal analysis on the biofilms as well as on the corresponding purified matrices. We showed that biofilms mechanical properties depending on the pEtN-cellulose to curli bacteria ratio are characteristic of a composite behavior of the ECM. Biofilm properties and ECM organization of the reference *E. coli* biofilms appeared to be mostly recovered in co-seeded biofilms with a 50:50 bacteria ratio. However, microscopic and spectroscopic investigations at the fiber scale revealed that the combination of curli fibers and pEtN-cellulose forms a product that is structurally different of its components, and that the hygroscopic properties of this product are different if the two components are produced and assembled by different (50:50) or by the same bacteria (AR3110).

The knowledge collected in this work will strengthen the promising use of biofilms as tunable materials and thereby benefit the synthesis of new bio-sourced composite materials,^12^ with applications as adhesives, textiles, filter membranes and bioinks to name a few.

## Results and Discussion

### 1. Contribution of ECM components pEtN-cellulose and curli to biofilm mechanical properties

To study the influence of the ECM components of *E. coli* biofilms, we mixed in different ratios and co-seeded bacteria producing only pEtN-cellulose (AP329) with bacteria producing only amyloid curli fibers (W3110) as the main extracellular component in their biofilm (**Figure 1a, Figure S1-2**). As reference, we seeded bacteria producing both main components pEtN-cellulose and amyloid curli fibers (AR3110).

**Figure 1.**
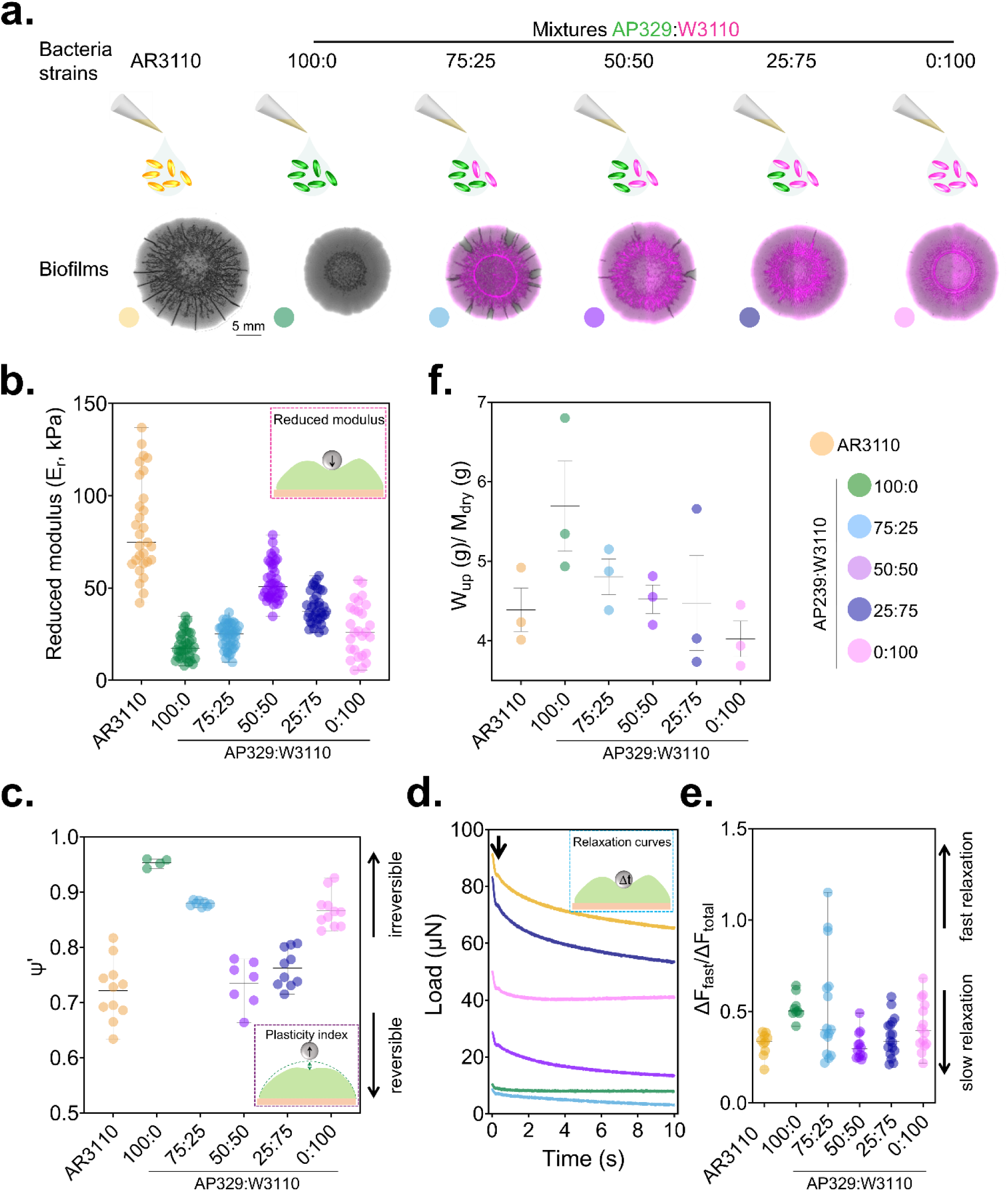
Mechanical characterization of the different strains of *E. coli* biofilms with micro-indentation. (a) Morphological features of the different biofilms according to the initial seeding composition of bacteria suspension. W3110 bacteria (curli producing bacteria) are m-cherry labelled to see the bacteria distribution in the biofilms. Independent experiments were done in 3-4 biofilms. (b) Averaged reduced elastic modulus performed on biofilm surfaces (10 individual measurements per biofilm, having 3 independent biofilms per strain). (c) Apparent plasticity index (ψ’) describes the capacity of the biofilm to deform irreversibly when a load is applied (Figure S4). (d) Representative relaxation curves of each strain during the holding time at a maximum penetration depth of 20 μm. The arrow shows the time limit (205 ms) chosen to calculate ΔF_fast_.^15^ (e) Force relaxation ΔF_fast_/ΔF_total_ derived from the relaxation curves. Mechanical data show minimum to maximum distribution bars with the median. (f) Water uptake per gram of dry biofilm calculated as W_up_ = (m_rewet_ – m_dry_)/_mdry_. Except for (a), data obtained correspond to indentation curves with a maximum depth of 20 µm. N = 3 different biofilms. The statistical analysis was done with Man Whitney U test (See Table S1- S4).

The growth rate of mixed bacteria and the projected area of the co-seeded biofilms were compared against AP329 (producing only pEtN-cellulose), W3110 (only curli) and AR3110 (pEtN-cellulose and curli) biofilms. No significant differences were observed in their bacterial growth curve (**Figure S1a**) nor in the initial phase of the biofilm growth (**Figure S1b**). The growth kinetics of the biofilms with different ECM composition only differed after the first 24 h of growth, i.e. after the onset of ECM production.^11^ Although there were significant differences between the projected areas of the AR3110 biofilms and those rich in pEtN-cellulose after 5 days of growth, no general trend was observed (**Figure S2b**). Fluorescent-tagged W3110 bacteria (mCherry) were co-seeded with AP329 non-fluorescent bacteria to observe whether there were areas in the biofilms where one of the bacteria types dominated (segregation) (**Figure 1a** and **Figure S1c**). Indeed, the top view of biofilms suggests higher segregation areas in the outer part of the biofilms initially containing more AP329 bacteria (**Figure 1a**). As the content of AP329 bacteria decreased in the biofilm, so did the number of segregation areas. Nevertheless, cross-sections of the biofilms showed that the bacteria were distributed evenly across the height of the biofilm (**Figure S1c**). What appears as a contradiction, could be due to the distribution of the bacteria in the proximity of the wrinkles and the limitations of the top-view imaging, where the projection of flat regions yields less signal than wrinkled regions. The more detailed method of cross-section imaging suggests co-seeding different mutants does not result in growth constrains nor in matrix segregation. The morphology of the different biofilms also appeared to vary with the initial seeding composition (**Figure 1a**, **S1 and S2b**). AP329 biofilms (pEtN-cellulose only) showed no visible wrinkling pattern, whereas W3110 biofilms (curli only) showed small radial wrinkles in the center of the biofilm. The biofilms formed with both components (pEtN-cellulose and curli) showed different patterns consisting in a combination of concentric and radial wrinkles (**Figure 1a**). While these patterns were observed in previous work,^7,14,15^ here we found that bacteria naturally producing both pEtN-cellulose and curli (AR3110) rendered biofilms with more defined wrinkling patterns compared to their co-seeded counterparts.

We then studied the contribution of the pEtN-cellulose and curli to the *E. coli* biofilms mechanical properties through micro-indentation experiments performed on intact biofilms on their agar substrate.^15^ To prevent a change in hydration state of the biofilms during storage, these measurements were done immediately after taking the plates from the incubator. Two different experimental set-ups were used whereby the indentation tip was either immediately retracted after reaching the maximum indentation depth, or held for 10s to allow force relaxation before retraction (**Figure S3-4**). On each load-displacement curve, considering the size of the tip (conospherical tip, r=50 µm) and the thickness of the biofilms (80-100 µm), we used the first 10 μm of the loading portion to calculate a reduced elastic modulus (E_r_) representative of the biofilm rigidity (**Figure 1b** and **Table S1**). The biofilms with the highest elastic modulus values were from AR3110 bacteria (pEtN-cellulose and curli) (median value 75 kPa) and co-seeded 50:50 (51 kPa). AP329 biofilms (pEtN-cellulose only) were the softest (17 kPa), while W3110 biofilms (curli only) were slightly more rigid, ca. 26 kPa. To assess the irreversibility of the deformation applied to the biofilms upon indentation, we defined an apparent plasticity index (ψ’) which describes the ability of the material to remain deformed when load is applied and immediately retrieved (**Figure S3, Figure 1c**, and **Table S2**). AP329 biofilms (pEtN-cellulose only) were the most plastic biofilms with an index of 0.95. However, the biofilms with the lowest apparent plasticity values, i.e. with higher elastic properties, were from AR3110 (0.73), co-seeded AP329:W3110 50:50 (0.75), and co-seeded AP329:W3110 25:75 (0.76). Note that these biofilms all contain a substantial amount of curli amyloid fibers in their ECM, always in presence of pEtN-cellulose. Comparative indentation experiments performed at the surface of water confirmed that the tip-sample interactions observed are not the result of an excess of free water accumulated as a layer at the surface of the biofilms (**Figure S3-4**).

The capacity of biofilms to relax forces was further studied while holding the indentation at the maximum depth for 10 s before retracting (**Figure 1d and e**, and **Figure S4**). The relaxation curves show a change of slope around 205 ms (**Figure 1d**).^15^ The first portion of the curve corresponds to a fast relaxation and was proposed to result from a poroelastic material response mediated by water flow.^15^ The second part of the relaxation curve corresponds to a slower relaxation mechanism, e.g., a reorganization of the biopolymer mesh. Here we assessed how much the biofilm relaxed before 205 ms with respect to the total amount of force relaxed (ΔF_fast_/ΔF_total_) (**Figure 1e** and **Table S3**). Biofilms with pEtN-cellulose only dissipated around 50 % of the total force relaxed in this short time laps (fast relaxation), whereas in biofilms producing curli only (W3110) the initial force relaxation is 40 %. In contrast, AR3110 biofilms showed similar percentage (approximately 30 %) of the force relaxed in this period of time as the co-seeded 50:50 and 25:75 biofilms. The capacity of the biofilm to dissipate indentation force during the holding time was described by the apparent holding plasticity (ψ’_h_) (**Figure S4-5, and Table S4**). ψ’_h_ followed the opposite trend as observed for ψ’ (**Figure S5a, and Table S4**) and for ΔF_fast_/ΔF_total_, and a similar trend as the reduced modulus (**Figure 1b**).

Lastly, the adhesion force F_ad_ required for the indentation tip to completely detach from the biofilm was calculated from load-displacement curves with a maximum indentation depth around 20 µm (**Figure S5b**, and **Table S5**). The strongest adhesion forces were measured on AR3110 and W3110 (curli only) biofilms (-16 and -26 µN, respectively), while AP329 biofilms (pEtN-cellulose only) presented lower adhesion (-11 µN). No significant differences were found among the co-seeded biofilms (AP329:W3110 75:25 -10 µN, AP329:W3110 50:50 -10.5 µN, AP329:W3110 25:75 -11 µN).

To assess the role of water in the contribution of *E. coli* ECM in biofilm mechanical properties, we measured biofilm wet mass, water uptake, and the osmotic gradients existing between the biofilms and their agar substrate. All biofilms were found to have a wet mass around 20 mg (**Figure S2c**). Yet, when biofilms were dried and rehydrated, a trend was observed where the higher the ratio of pEtN-cellulose producer seeded in the biofilm, the higher the water uptake (W_up_) per dry mass of the biofilm (**Figure 1f**). The AP329 (pEtN-cellulose only) biofilm showed a W_up_/M_dry_ ratio of 5.7 ± 0.98 gram of water uptake per gram of dry mass, while the W3110 (curli only) biofilm showed the lowest ratio of 4 ± 0.4. The osmotic pressure gradients (Δ∏) inform about the influx of nutrient-rich water from the agar substrate into the biofilm (**Figure S2d**). AR3110 biofilms showed the highest Δ∏ of ∼ 1.5 MPa, while no significant differences were found among the Δ∏ of the other biofilms (∼ 0.7 MPa).

The results obtained through this mechanical characterization of biofilms with different ECM compositions agree with the literature describing AR3110 *E. coli* biofilms as hydrogel composites made of stiff curli amyloid fibers and soft pEtN-cellulose carbohydrates assembled in a material that is greater than the sum of its parts (**Figure 1**).^15,21,22^ Moreover, the progressive introduction of each ECM component in the biofilm by seeding in different ratio bacteria exclusively producing pEtN-cellulose (AP329) or curli (W3110) demonstrated that a given balance of both components is necessary to recover the macroscopic mechanical properties of the reference biofilm (AR3110), and that this balance is partially achieved by co-seeding AP329 and W3110 in equal proportions (50:50).

pEtN-cellulose on its own is relatively less rigid compared to curli amyloid,^13,15,23^ and this ability is reflected in the respective biofilm reduced moduli (**Figure 1b**). Yet, understanding biofilm plastic behavior upon unloading requires to consider that pEtN-cellulose is proposed to be amorphous and zwitterionic, which promotes its interaction with water^24^ and explains the increasing water uptake measured in biofilms with increasing pEtN-cellulose content (**Figure S2d**). The high affinity of pEtN-cellulose with water can also explain that its presence reduces biofilm adhesion to the indentation tip (**Figure S5b**). The apparent plasticity calculated after short-time indentation (ψ’, **Figure 1c**) indicates that biofilms containing more pEtN-cellulose are more prone to an irreversible deformation, despite undergoing lower loads (**Figure S3**), potentially due to rapid changes in water distribution in the ECM. This is supported by the large ΔF_fast_/ΔF_total_ measured while holding the tip at maximum indentation (**Figure 1e**), indicating that the fast relaxation is most probably mediated by water.^15^ Moreover, the lower apparent holding plasticity calculated during 10 s at the maximum indentation depth (ψ’_h_, **Figure S5a**) indicates that the same biofilms are more prone to shape recovery, suggesting that the pEtN-cellulose polymer does not reorganize much during this holding time. We thus propose that indentation rapidly expels water initially adsorbed on the soft pEtN-cellulose fibers, which also greatly deforms as the tip reaches its maximum depth (**Figure 1e**), and that holding the tip favors water re-adsorption (swelling) by the highly charged groups of the pEtN-cellulose upon unloading (sponge-like effect). The ECM would then adopt a similar arrangement as before indentation.

In contrast, stiffer curli-containing biofilms appeared to be slightly less prone to irreversible deformation especially when pEtN-cellulose is also in the ECM (**Figure 1c**). Water is still expected to play a role, however the relatively slower force relaxation measured in such biofilms seems dominated by ECM re-organization (**Figure 1e**). While curli fibers are rigid and brittle,^14,24,25^ they may break under loading as reflected in the relatively high apparent plasticity index of W3110 biofilms (curli only) (**Figure 1c**). Yet, the sole presence of pEtN-cellulose in curli-containing biofilms, even in small proportion, was sufficient to decrease their apparent plasticity and slowdown force relaxation most probably through matrix reorganization (**Figure 1e**).^21,23^ Since the evolution of biofilm elastic and relaxation properties as a function of pEtN-cellulose content is not linear and do not scale with biofilm water uptake (**Figure 1f**), we propose that interactions between curli and pEtN-cellulose at lower scales play a crucial role in creating a synergy between their respective properties and leading to the observed composite behavior.^15^

The determining role of interactions between curli and pEtN-cellulose is supported by the comparison of biofilms from co-seeded bacteria (AP329:W3110) with biofilms from the reference bacteria strain producing both components (AR3110). AR3110 biofilms are not only significantly larger (**Figure 1f**) and more rigid (**Figure 1b)** than the other biofilms, but they also induce higher osmotic gradients with the agar (**Figure S2d**) and show relatively high adhesion (**Figure S5b**).^21^ Explaining the observed differences required to study the role and the nature of the interactions between the ECM components and thus characterize the different biofilms at lower length scales.

### 2. Mechanical roles of pEtN-cellulose and curli for the biofilm morphology

To assess biofilm architecture as a function of ECM composition and understand its role in the macroscopic properties described above, structural characterizations of the ECM were then performed within the different co-seeded and reference biofilms.

Direct Red 23 (also known as Pontamine Fast Scarlett 4b) has affinity for pEtN-cellulose and curli amyloid fibers, which makes it a suitable dye to visualize ECM in biofilm cross-sections by confocal microscopy (**Figure 2**, **Table 1** and **Figure S6a**).^7,18^ Imaging cross-sections of the different biofilms first enabled to compare their respective thicknesses. Biofilms from W3110 bacteria (pEtN-cellulose-:curli+) were the thickest, while AP329 biofilms (pEtN-cellulose+:curli-) were the thinnest (**Table 1**) (**Figure S6a**). The co-seeded biofilms containing both pEtN-cellulose and curli presented increasing thickness values, as the presence of curli producing bacteria increased (**Table 1**). AR3110 biofilms naturally producing both components had an averaged thickness of 77 ± 8 µm.

**Figure 2.**
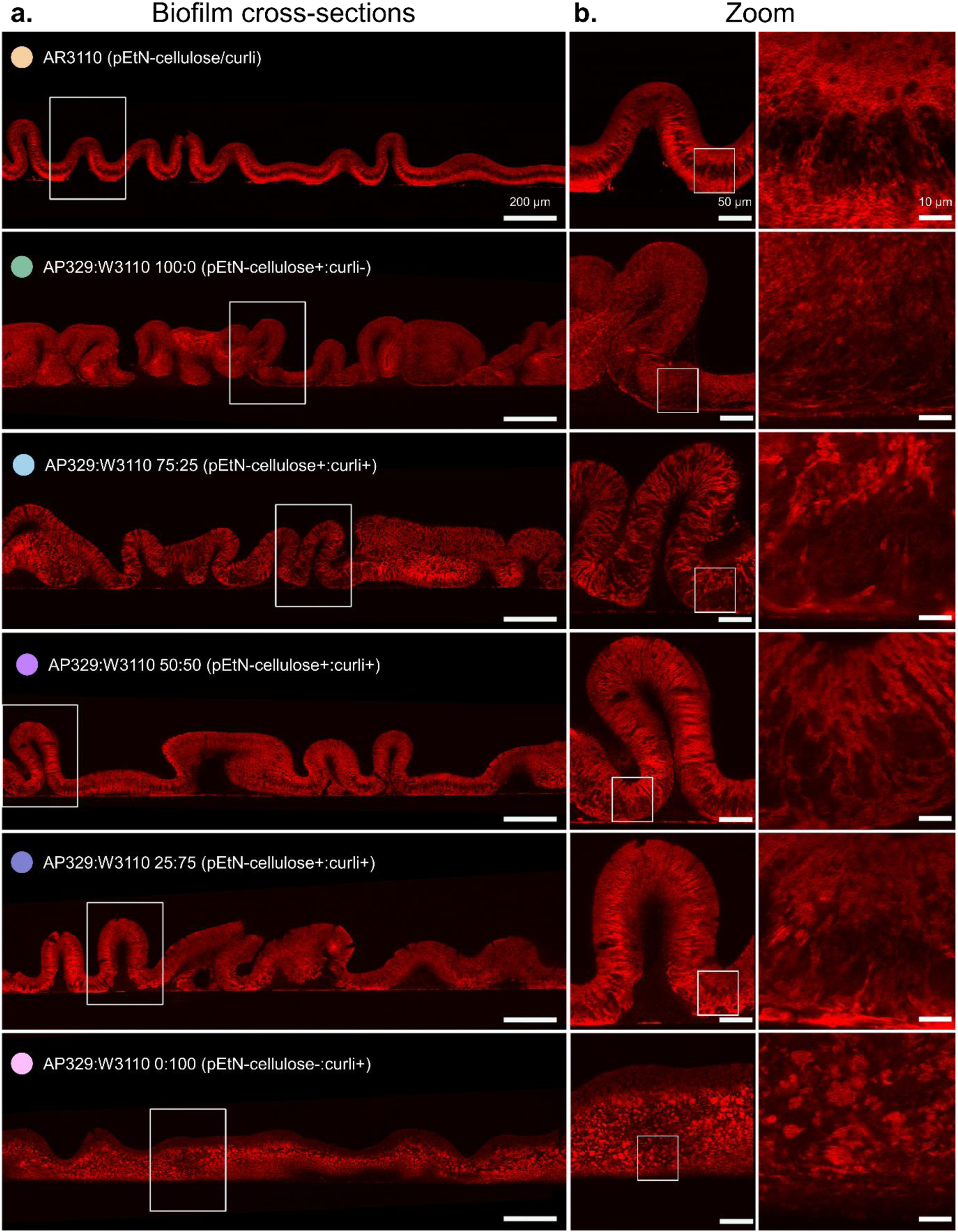
Microarchitecture of extracellular matrix of the different biofilms in the study. (a) Confocal microscopies of cross-section of the biofilms stained with Direct Red 23. (b) Zoom of cross-sections of the different strains of biofilms in (a). The respective area of the zoom of the cross-section are indicated by squares of each area. N=6

**Table 1.**
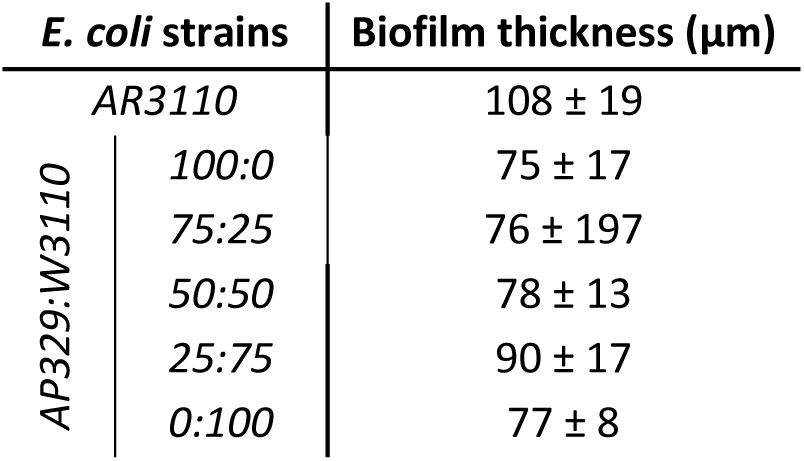
Biofilm thickness. The average values in the table are extracted from. **Figure 2 and Figure S7a. (N= 6)**

Biofilm cross-sections also revealed the wrinkling pattern of the biofilms along the third dimension. Biofilms containing only curli (W3110) showed much less wrinkles than the others (**Figure S6b,** and **Table S6**).^14,15^ Note that AP329 biofilms did have small and densely packed wrinkles that were not visible on top-view images (**Figure 1a**). The presence of pEtN-cellulose in the ECM was necessary to achieve the characteristic well-defined wrinkling pattern observed in the reference AR3110 biofilms (pEtN-cellulose and curli) (**Figure 2**). The cross-sections from the co-seeded biofilms indicated that the ratio of pEtN-cellulose to curli in the ECM is not as important as their sole presence in order to obtain a wrinkling pattern similar to AR3110 biofilms. Yet, the wrinkling pattern increased with the content of pEtN-cellulose producing bacteria (AP329) in the biofilm (**Figure S6b**).

Finally, the fluorescence images of biofilm cross-sections with stained ECM showed that the spatial arrangement of the fibrous matrix varies depending on the presence/absence of pEtN-cellulose and curli (**Figure 2** and **Figure S6a**).^26^ When both pEtN-cellulose and curli were present in the AR3110 biofilm, cross-sections revealed a structured spatial arrangement in which fibers were clearly oriented perpendicular to the direction of the biofilm. There were clear defined areas of orientation of the fibers or density (depicted qualitatively by fluorescence intensity, **Figure S6a**).^7^ The top and bottom parts of the biofilm depicted a dense network (brick-like) of fibers, while the middle portion showed fibers arranged in a vertical way, as described in previous work.^13^ This pattern was partially retrieved by the co-seeded biofilms with pEtN-cellulose and curli amyloid fibers to different extents. In contrast, the biofilms from AP329 bacteria presented a uniform mesh of pEtN-cellulose. Finally, W3110 biofilms presented a patch-like spatial arrangement of the curli fibrous matrix, with aggregates of about 10 µm size.^18^

Biofilm morphology can be interpreted in light of these mechanical data (**Figure 2**).^27^ AP329 shows wrinkles with higher local curvature than AR3110, some of them even falling over to form creases. This is in line with a lower bending rigidity of the biofilm, probably due to the absence of curli. Indeed, the measured modulus of AP329 is about three times lower than for AR3110 that secretes both curli and pEtN-cellulose (**Figure 1b**). In contrast, W3110 lacking cellulose shows no wrinkles, although the biofilm rigidity is about as low as for AP329 (**Figure 1b**). However, this biofilm has a swelling ratio about 40 % lower than AP329 (**Figure S2d**). This seems to indicate that enhanced water uptake by the presence of pEtN-cellulose is needed for the formation of wrinkles (since W3110 lacking cellulose does not form wrinkles) and that a stiffening of the matrix by the presence of curli is required to prevent the wrinkles from collapsing to creases (as seen in AP329 lacking curli). Mixtures of AP329 and W3110 recover most of the wrinkling pattern of AR3110, which indicates that the interaction of pEtN-cellulose and curli is essential for biofilm geometry by combining water uptake and rigidity.

To better assess the subtle differences between the AR3110 and co-seeded 50:50 ECMs, we used cryo-Focused-Ion Beam Scanning Electron Microscopy (FIBSEM) to image small volumes of the corresponding biofilms at higher resolution (**Figure 3a**). For this, we first defined three regions according to their morphological differences and bacterial activity: a central zone forming a disk reminiscent of the initial seeding drop and where a dense ECM often supports the formation of wrinkles, an outer zone appearing as a flat rim where active bacteria proliferate and start to produce new ECM, and an intermediate zone in-between.^11,13,28^ We then imaged a volume of each zone in both biofilms from AR3110 bacteria and from co-seeded bacteria with a ratio of 50:50 (**Figure 3a**). We observed differences between biofilms in the organization of their ECM.

**Figure 3.**
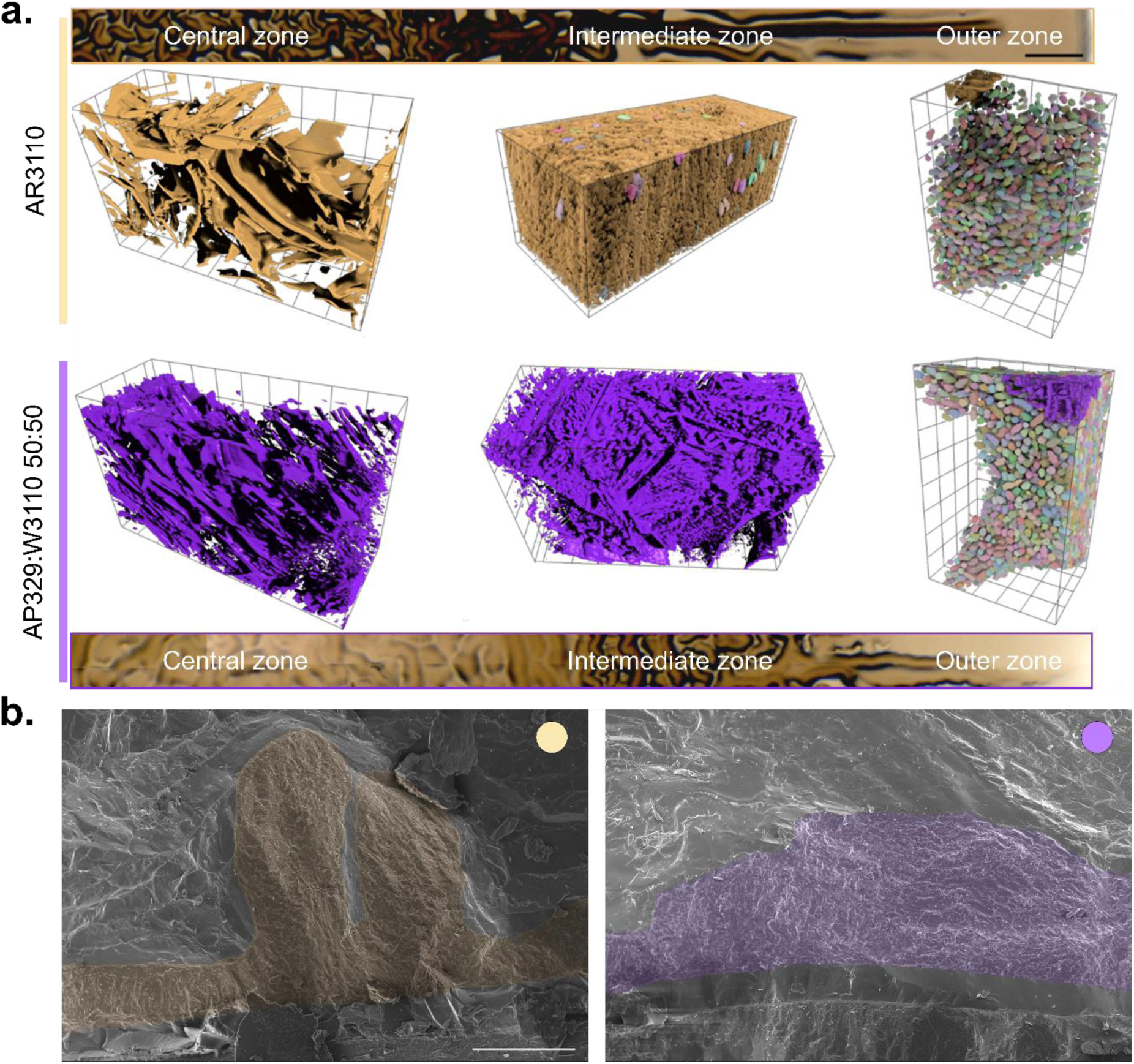
Biofilm ultrastructure. (a) 3D perspective rendering of the segmented different regions of different *E. coli* biofilms: AR3110 and AP329:W3110 50:50 acquired in cryo-FIBSEM. Top sketch depicts where the imaged regions belong to in the biofilm (center, intermediate and outside zone). Grid size= 3 µm (b) cryo section ESEM images of *E. coli* biofilms: AR3110 and AP329:W3110 50:50. Images were acquired at 800x magnification. Scale bar= 100 µm.

Altogether, the samples taken from each zone of the biofilm depicted the transition from mature to new ECM as described by Klauck et al.^28^ The central zone of the AR3110 biofilm showed ECM fibers with an organized spatial arrangement following a sheath-like structure. Qualitatively, these ECM fibers were denser and more organized than the ECM fibers found in the co-seeded biofilm. The outer zone of both biofilms presented similarly high bacteria and low ECM concentrations, in agreement with previous descriptions.^13^ In both cases, the intermediate zone appeared to be denser and with a more compact arrangement of the biofilm ECM, potentially due to high metabolic activity in this region.^28^ Comparing both types of biofilms, we observed that the co-seeded biofilm presented again a more disordered ECM arrangement compared to AR3110. In this area, we found a few AR3110 bacteria embedded in their fiber mesh, but not in the co-seeded. Note that the absence of bacteria in some of the volumes acquired by cryo-FIBSEM imaging could be either due to the choice of the location or to a specific contrast locally yielded by the fibers which impairs their simultaneous visualization together with bacteria. Alongside, freeze fracture images of the intermediate zone imaged by cryo-ESEM showed that the AR3110 biofilm matrix is spatially more organized than the mixed biofilm matrix (**Figure 3b**). While the AR3110 ECM fibers seemed to have a preferred vertical orientation (from top to bottom) all along the biofilm, the ECM fibers belonging to the co-seeded biofilms appeared to have no orientation at all.

The microstructural differences found between AR3110 and co-seeded 50:50 ECMs (**Figure 3**), especially in the degree of organization of the fibrous matrix, demonstrate the importance of the conditions of interactions between pEtN-cellulose and curli. Indeed, in AR3110 biofilms, both pEtN-cellulose and curli are simultaneously produced by the same bacteria (*E. coli* AR3110) in a co-dependent manner,^29^ which implies that the two components interact in a short range, shortly after exiting the bacterium. In contrast, the co-seeded biofilm is composed of the two different *E. coli* mutants AP329 (producing only pEtN-cellulose) and W3110 (producing only curli amyloid fiber), so that pEtN-cellulose and curli can only interact once they find each other in the extracellular space. We thus propose that the co-secretion of pEtN-cellulose and curli in AR3110 biofilms favors ECM organization at the microscopic scale by promoting their interactions right after secretion.

### 3. Characterization of pEtN-cellulose, curli and their assembly in *E. coli* ECM

Mechanical and structural characterizations showed that biofilms co-seeded from AP329:W3110 50:50 better recovered the properties of reference biofilms (AR3110) than the other co-seeded biofilms, and the remaining differences were proposed to derive from differences in pEtN-cellulose and curli fiber interactions. Therefore, we investigated whether the structure of purified ECM components could explain these differences and similarities. Following the same extraction method used for curli amyloid fibers,^30,31^ we purified the matrix from biofilms obtained from AR3110 (pEtN-cellulose and curli), AP329 (pEtN-cellulose only), W3110 (curli only) and co-seeded 50:50 (pEtN-cellulose and curli) (**Figure 4a**). Interestingly, despite various attempts, it was possible to purify but not separate the curli and pEtN-cellulose fibers from the biofilms containing both components. Depending on the co-seeding ratio, the purified fibers were thus either pEtN-cellulose, curli, or a mixture of curli and pEtN-cellulose.

**Figure 4.**
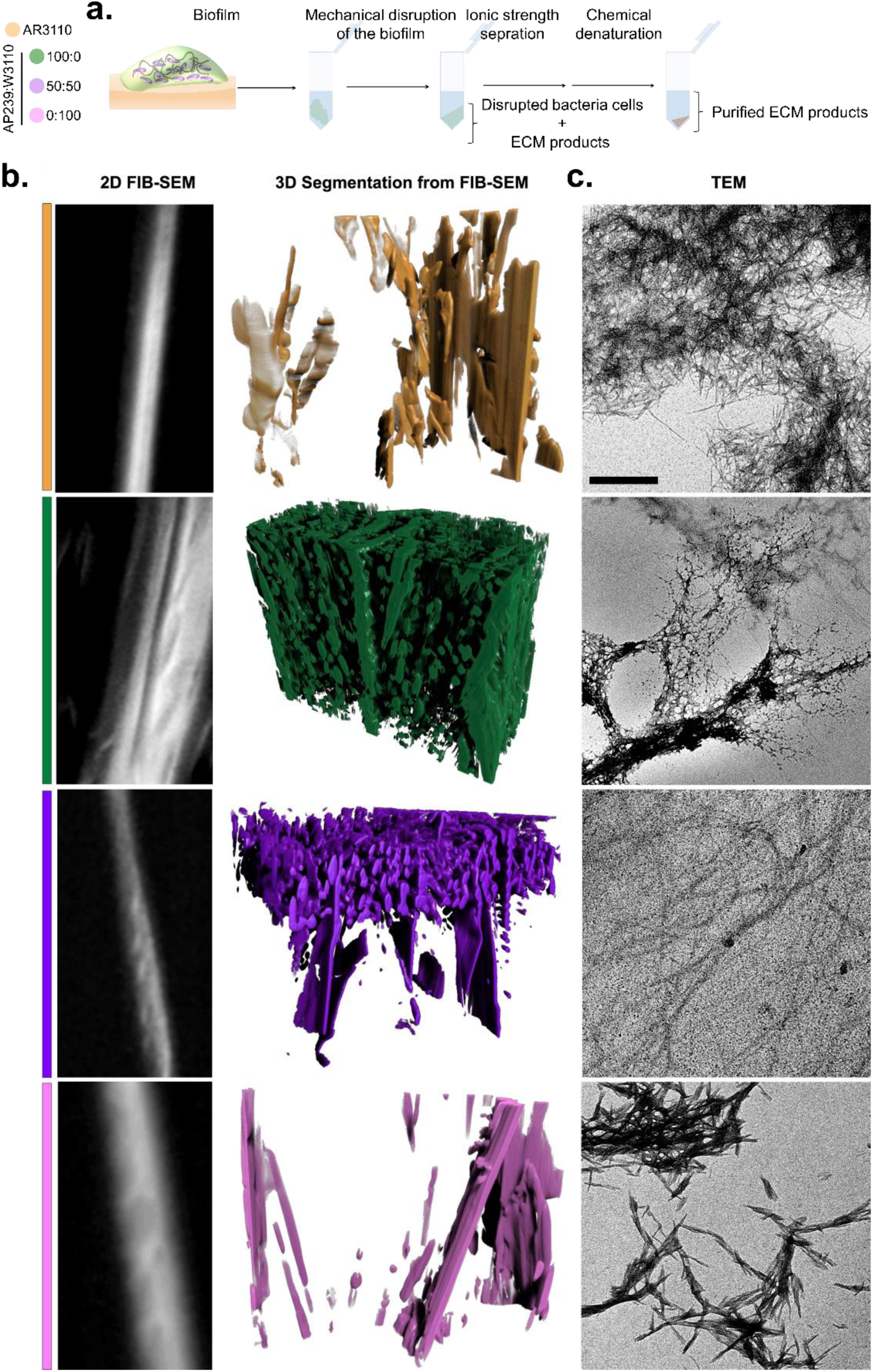
Electron microscopy images and segmentation of purified products from the *E. coli* biofilms. (a) Scheme of the extraction process for the ECM components. (b) Representative image of the purified fibers extracted from the cryo-FIBSEM image stack (left column), and 3D rendering of each type of purified fibers reconstructed from cryo-FIBSEM stack (right column). (c) Transmission electronic microscopy images of the purified fibers. Data come from 4-5 independent biofilm cultures for each condition tested. For all electron microscopies the scale bar = 500 nm.

Cryo-FIBSEM was used to image the purified fibers in three dimensions (**Figure 4b**). We observed different types of spatial organizations of the fibers, depending on their composition. Purified pEtN-cellulose (AP329) appeared as a dense and disordered assembly of sheath-like structures separated by some globular aggregates. In contrast, purified curli amyloid fibers (W3110) appeared as dispersed sheath with a needle-like morphology. AR3110 purified fibers displayed a similar organization, but less ordered than in pure curli (W3110) and less dense than in AP329 fibers. The purified fibers from the co-seeded biofilms showed a mixture in both spatial order and morphology between the AP329 and W3110 purified fibers. A close-up from the cryo-FIBSEM stack of each purified ECM revealed a smoother and uniform texture of the AR3110 purified fiber compared to the fibers obtained from co-seeded biofilms. The latter ones presented a more porous structure with twists. These twists were also observed in the AP329 fibers, while the purified curli amyloid fibers showed a general smoother surface, but some twists along the fiber were still visible.

When observed with transmission electron microscopy (TEM), we realized the fibers observed in cryo-FIBSEM are in fact bundles of each fiber type. Thus, TEM measurements revealed details in the morphology of the fibers, and confirmed the results observed (**Figure 4c**).^17,18,31^ The pEtN-cellulose matrix fibers purified from AP329 biofilms organized into a mixture of bundles and sheets as observed in other types of bacterial cellulose,^32–34^ whereas curli amyloid fibers from W3110 biofilms were short and needle-like.^17,31,35^ ECM purified from AR3110 biofilms (containing both pEtN-cellulose and curli) also showed needle-like fibers but longer and thinner than in pure curli. Finally, the fibers from co-seeded biofilms were even longer and slightly bent reminiscent of a mixed organization between the W3110 and AP329 purified products.

To assess the interactions between pEtN-cellulose and curli, we then applied biophysical characterization techniques well established on amyloid fibers to the different purification products (**Figure 5**). Thioflavin T (ThioT) is a fluorescent probe used to identify β-sheet-rich structures such as protein amyloid fibers.^17^ The intensity of this probe is related to the packing, length and amount of these structures.^17,36,37^ Other than the purified pEtN-cellulose from AP329 biofilms, all purified fibers presented ThioT emission (**Figure 5a**), confirming that the extracted fibers from *E. coli* AR3110 biofilms (natural producers of pEtN-cellulose and curli) and from co-seeded biofilms contained amyloid-like fibers in their composition. As expected, the biofilms from *E. coli* W3110 (curli amyloid fibers only) showed the highest ThioT emission. The differences in intensity suggest differences in fiber structure in terms of β-sheet arrangement, or concentration.^17,38,39^

**Figure 5.**
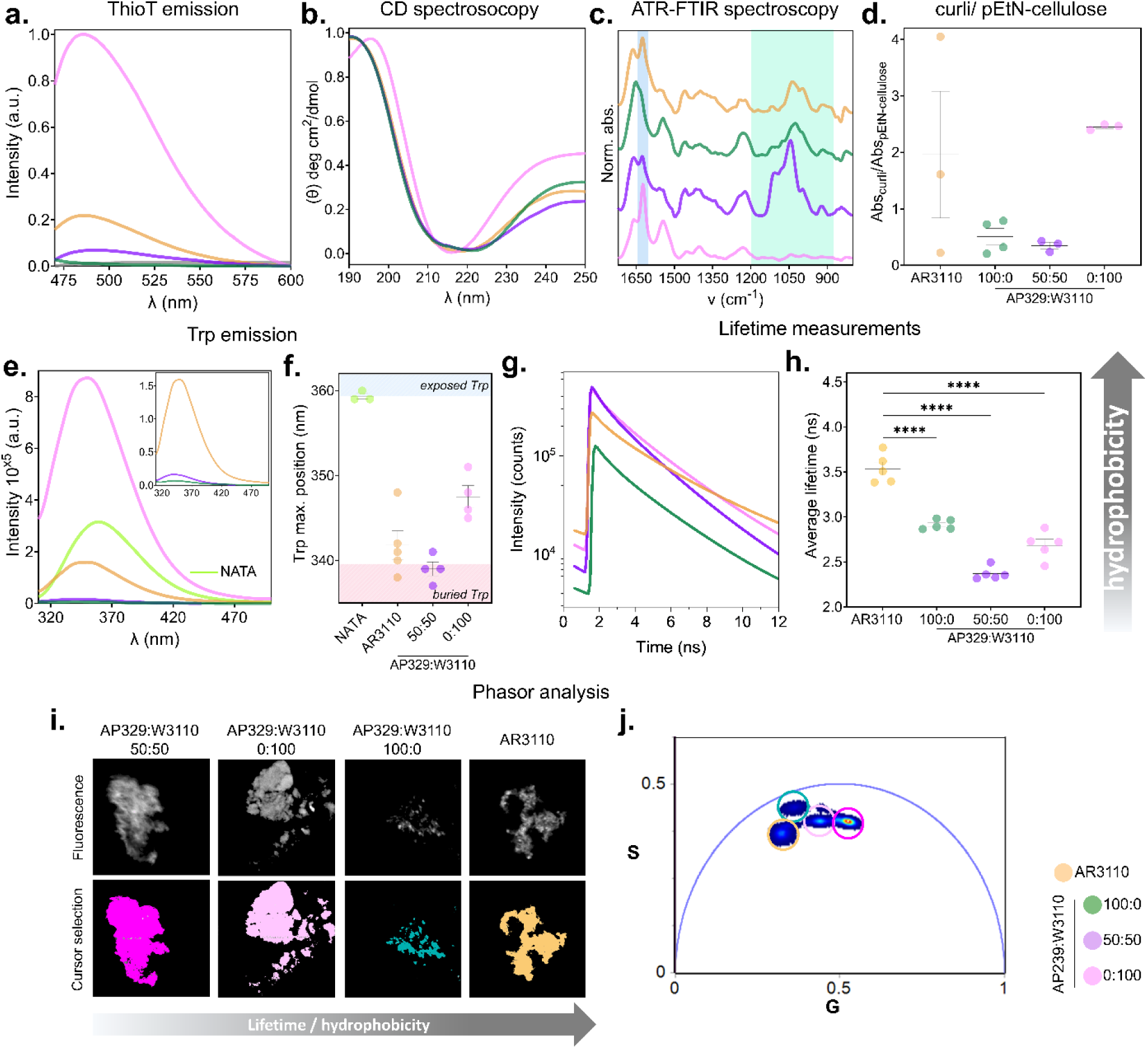
Characterization of the structure of purified curli fibers. (a) ThioT fluorescence emission spectra of the fibers after purification. (b) Circular dichroism (CD) spectroscopy of the purified fibers. (c) Representative Amide I’ region of an ATR-FTIR spectrum for the purified fibers. The protein and carbohydrate regions used in the analysis are highlighted in blue and green, respectively. (d) Estimation of the composition of the purified fibers extracted from the curli/pEtN-cellulose ratio from panel (c). The statistical analysis was done with One-way ANOVA (p<0.0001, **** | p<0.001, *** | p<0.01, ** | p<0.05, * | ns = non-significant), post-test used was Tukey’s test to compare each fiber against every fiber (**Table S7**). (e) Intrinsic emission spectra of the Trp population of the purified fibers containing curli. The spectrum of soluble Trp (NATA) in buffer is represented as reference for the maximum exposure possible of the Trp to the surface (λexc = 280 nm). (f) Position of the emission peak of the intrinsic fluorescence of the fibers in (a). The shadowed areas in the main plot indicate these exposure extremes; exposed Trp (blue) and buried Trp (red). N=3-4 independent biofilm cultures for each fiber tested. (g) Lifetime measurements of Nile Red bound to the different purified fibers. (h) Average lifetime values for Nile Red in the different systems. Lifetime curves were fitted with three components (see **Table S8**) and the average values weighted by intensity are plotted. n=3, statistical analysis was done with One-way ANOVA (p<0.0001, **** | p<0.001, *** | p<0.01, ** | p<0.05, * | ns = non-significant), post-test used was Tukey’s test to compare each fiber against every fiber (i) Confocal and FLIM images of the purified fibers for each condition bound to Nile Red.^18^ (j) Phasor plot of the FLIM images for each sample. The pixel clouds show colors from blue to red according to the pixel density. The S and G axis represent the imaginary and real components of the Fourier transform (see methods for details). The reciprocity principle allows to select the different pixel clouds with cursors that will color the correspondent pixels in the images. N=3 independent purification batches of fibers were used in each experiment.

We used circular dichroism (CD) and ATR-FTIR spectroscopy to analyze the secondary structure of the purified fibers from each biofilm (**Figure 5b-c**, and **Figure S7**). CD spectroscopy identified further differences in the structure of the fibers (**Figure 5b**). W3110 curli amyloid fibers and AP329 pEtN-cellulose were used as extreme references for comparison with the mixed purified products. The CD spectra of the fibers purified from AR3110 and co-seeded (50:50) biofilms were similar, but different from the AP329 pEtN-cellulose, and W3110 curli spectrum (**Figure 5b**). Overlapping signals from pure curli and pure pEtN-cellulose in the mixed fibers spectra prevented from further analysis.

Using ATR-FTIR spectroscopy, we observed that both the absorption spectra of the purified fibers from AR3110 and the co-seeded biofilms (50:50), as well as their second derivative are different compared to the pure fibers (pEtN-cellulose alone or curli alone) (**Figure 5c**, and **Figure S7**). The spectra of these fibers showed similarities with those of pure curli amyloid fibers in the amide I region (1600 – 1700 cm^-1^), with a characteristic peak around 1622 cm^-1^, signature of the presence β-sheet structure.^17^ In contrast, the pure pEtN-cellulose spectrum presented a peak at 1025 cm^-1^ (in the fingerprint region of polysaccharide 1200 – 900 cm^-1^) corresponding to the stretching vibration C−O of secondary alcohols.^25^ The Amide I’ region in the curli spectrum (1600 – 1700 cm^-1^) represents the stretching vibrations of the C=O and C-N groups.^40^ Within this region, we focused on the 1622 cm^-1^ maximum, characteristic of intermolecular β-sheets (**Figure 5c**).^41^ Both spectra from AR3110 and co-seeded fibers presented a shift by ≈10 cm^-1^ for the peak around 1025 cm^-1^ and ≈5 cm^-1^ for the peak around 1622 cm^-1^. These shifts suggest an interaction between pEtN-cellulose and curli in the composite fibers. To estimate semi-quantitatively the composition of the purified products, we calculated the ratio between the signature peak for β-sheet of the protein amyloid fibers (1622 cm^-1^) and the region between 1200 – 900 cm^-1^ characterizing pEtN-cellulose (**Figure 5d**, and **Table S7**). The higher the ratio, the higher the content of pEtN-cellulose in the purified ECM. Despite the high dispersion, the composition of the purified ECM from AR3110 biofilms was different from the fiber composition of purified ECM from the co-seeded biofilms. When produced by the same bacteria (AR3110), the curli/pEtN-cellulose average ratio was ≈2. When the fibers were produced by different bacteria in the same biofilm (co-seeded), the curli/pEtN-cellulose average ratio was ≈0.5, which suggests a lower curli content and/or a higher pEtN-cellulose content in this type of fibers. The dispersion of the ratio obtained from AR3110 ECM, could be due to the estimation method, to the differences in the ATR-FTIR spectra obtained, or to natural fiber polymorphism or composition during the polymerization of the fibers. Bradford’s test was used to further estimate fibers protein content (**Figure S8**). The protein concentration in the purified fibers from co-seeded biofilms 50:50 was higher than in the purified fibers from AR3110 biofilms. While the estimation by ATR-FTIR reports the ratio between curli and pEtN-cellulose in the fiber composition, the protein estimation by Bradford’s reports the absolute concentration of protein in the fibers.

We then studied the intrinsic fluorescence of the purified ECM using the tryptophan (Trp) present in the CsgA curli monomer as a probe (**Figure 5e-f**).^18^ The Trp residue fluorescence is sensitive to its direct nano-environment: its spectrum shifts toward shorter wavelengths as it surroundings becomes more hydrophobic.^40,42^ We used soluble Trp (NATA) as a reference for the spectrum of a Trp with the highest possible exposure (i.e. the most hydrophilic environment).^17^ Since pEtN-cellulose does not have a Trp in its composition, we focused our comparison on the mixed samples and the purified curli fibers obtained from W3110 (curli only) biofilms (**Figure 5e, inset**). The position of maximum Trp emission indicated that Trp in the purified W3110 curli fibers were much more exposed than the Trp in purified fibers from AR3110 and co-seeded biofilms (**Figure 5f**). Although similar, the Trp in the purified fibers of co-seeded biofilms were slightly less exposed to the solvent than the Trp in the purified AR3110 products.

As another approach to understand the nano-environment of the different ECM components, we used an external dye, Nile Red (NR). NR, similar to Trp, is a solvatochromic probe that changes its spectrum and lifetime in response to its environment. It has the advantage that it can be added externally to the sample, and it binds into hydrophobic pockets present in fibers, allowing to distinguish the polymorphism.^17,39^ We used fluorescence lifetime imaging (FLIM) to measure the lifetime of NR and to study the hydrophobicity of the purified fibers (**Figure 5g-i**).^17,39^ The lifetime of NR increases when the hydrophobicity of its environment is higher (**Figure 5g**). The results show that the binding site from AR3110 biofilms purified fibers (pEtN-cellulose and curli) have a significantly higher hydrophobic character than the rest of purified ECM components.^17,43,44^ The NR bound to the purified fibers from the co-seeded biofilm presented the lowest hydrophobic character (**Figure 5h**).

While differences in the average NR lifetime values already suggest there are differences in the hydrophobicity of the fibers, combining FLIM with the phasor approach allows further interpretation of the results.^17,45^ The phasor analysis is a representation of the raw data in a polar plot that allows a direct interpretation of the changes occurring in the system without the assumption of any models a priori.^45^ **Figure 5i-j** show confocal microscopy images of fibers stained with NR and the corresponding phasor plot for the lifetime of the different purified fibers. In **Figure 5j** each colored cursor encircles the position of the pixels of a given lifetime of NR bound to the different fibers. Due to the properties of the Fourier space, if the mixed fibers from the AR3110 and co-seeded biofilms were perfect mixtures, the pixels corresponding to them should fall on an linear trajectory between the pure components, i.e. the pEtN-cellulose (green circle) and curli (pink circle) groups of pixels (**Figure 5j**).^46^ The fact that the purified fibers from the AR3110 and co-seeded biofilms fall outside the linear trajectory between the pure components, strongly suggests that these fibers are not only different from each other, but form a material which properties are not just the addition of the two components, but rather a new distinct composite material.

The existence of a curli/pEtN-cellulose composite behavior at the fiber level is supported by the differences observed between the pure fibers (ECM from AP329 or W3110 biofilms) and the mixed fibers (ECM from AR3110 and co-seeded 50:50 biofilms) using multiple analysis modalities (**Figures 4-5**). Indeed, assessing the composition of purified ECMs by ATR-FTIR and Bradford experiments indicated differences between the AR3110 and 50:50 fibers, thereby suggesting that the composition of protein and carbohydrates in the purified biofilm ECM fibers is influenced by the way the fibers were polymerized and assembled during biofilm growth (**Figure 5d and S7**). Moreover, differences in ThioT signal intensities suggest differences in fiber β-sheet arrangement or concentration (**Figure 5a**).^17,38,39^ The slight differences in their CD spectra and Trp positions also suggested differences in either their structure and/or the way the curli and pEtN-cellulose assembled at the fiber level (**Figure 5f-i**). The NR bound to the 50:50 fibers showed the least hydrophobic character of all fibers, while their ATR-FTIR spectra suggest interactions between the protein and carbohydrate portions of these fibers (**Figure 5c-j**, and **Figure S7**). The groups involved in the interaction between the pEtN-cellulose and curli portion of the fibers are the same for the AR3110 and 50:50 mixture: C-O of secondary alcohols^25^, and C=O and C-N groups^40^, respectively. This suggests that while the chemical groups of the pEtN-cellulose and curli interacting with each other might be the same, the resulting structures are different.

### 4. Model for the interactions between curli and pEtN-cellulose

The results of this work strongly suggest that the composite behavior of *E. coli* biofilms ECM containing both curli and pEtN-cellulose arises from molecular interactions happening either at the supra-bacterial scale in the case of co-seeded biofilms (different bacteria), or at sub-bacterial scale in the case of the reference strain AR3110 producing both components (hybrid composite material) (same bacteria). Two possible scenarios of interactions can be envisaged: 1) curli and pEtN-cellulose could assemble “in series”, with portions of pEtN-cellulose intercalating in-between csgA monomers as the curli fiber is assembled, or 2) curli and pEtN-cellulose could assemble “in parallel”, with curli and pEtN-cellulose fibers interacting along each other as they meet in the extracellular space. Both scenarios are not exclusive and could happen in AR3110 biofilms where both fibers are produced by the same bacteria (**Figure 6a**, yellow). In co-seeded biofilms however, the two components are likely to interact further away from the bacteria, i.e. as they are already assembled into fibers, which would then favor scenario 2 (**Figure 6a**, purple). The fact that the same groups are involved in the interactions in AR3110 and in 50:50 supports the dominance of scenario 2 in both cases (**Figure 5c and S7**). The interactions in co-seeded biofilms (50:50) would then be of same type but less dense than in AR3110, in agreement with the cryo-FIBSEM and TEM data (**Figure 3, 4b and 4c**). Such interactions may also stabilize each type of fiber during their formation, thereby leading to increased ECM fiber length and/or thickness (**Figure 4c**), which are in turn expected to influence biofilm structure and mechanical properties (**Figure 2 and 1)**. A study of the interaction between oligosaccharides containing the pEtN-group with synthetic peptides representative of csgA showed that tyrosine, glutamine, histidine and serine are amino acid affected by the presence of the oligomers.^20^ Interestingly, the few tyrosine entities are located rather at the extremities of the csgA monomer, whereas glutamine, histidine and serine are rather distributed across the peptide. This indicates that both scenarios are plausible to describe the emergence of the next hierarchical level.

**Figure 6.**
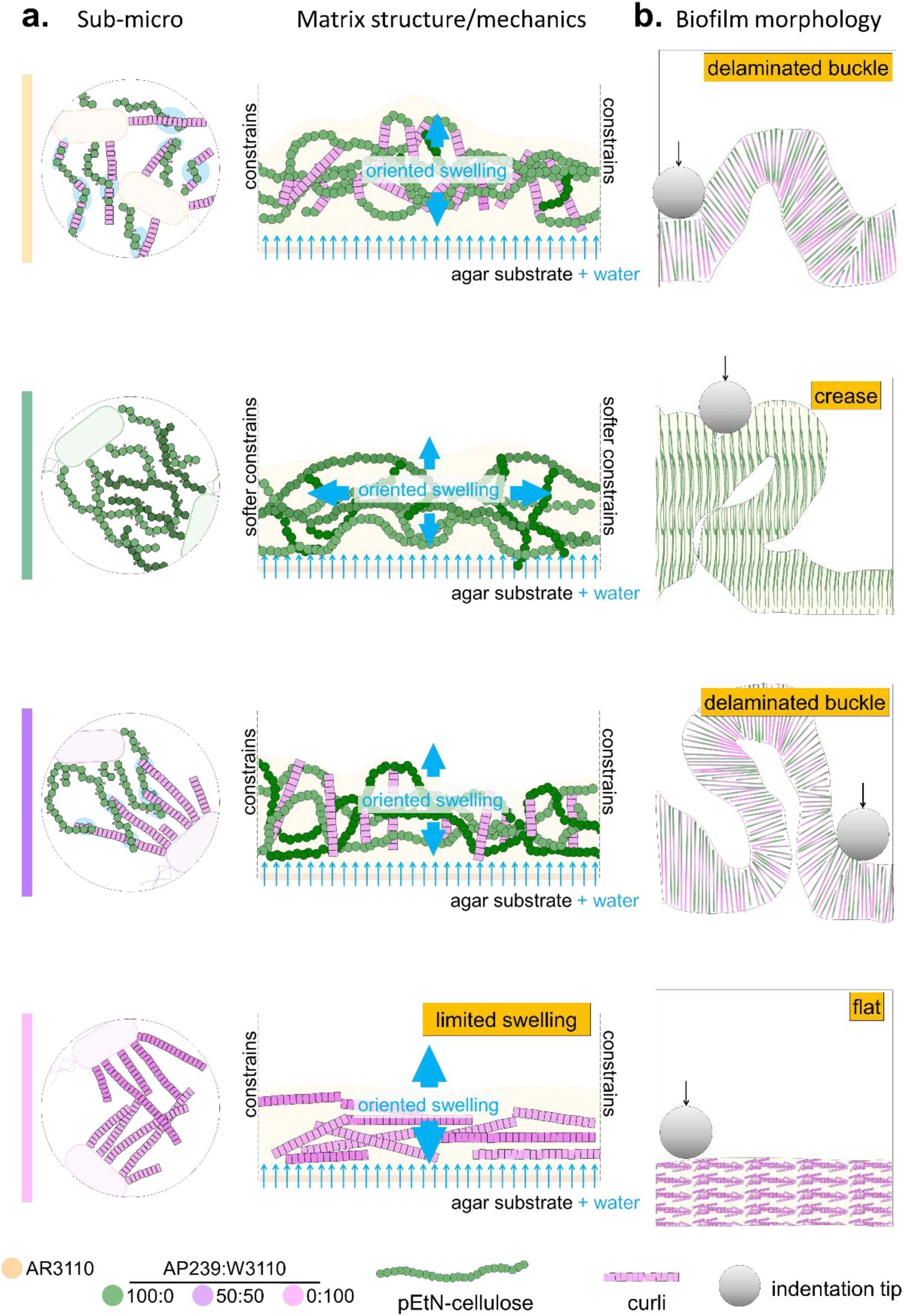
A model for the contributions of the ECM components to *E. coli* biofilms behavior. (a) Representation of a possible fiber assembly scenario at the bacterial scale. (b) Mechanical behavior of the curli / pEtN-cellulose composite at the mesoscopic scale. (c) Consequences of *E. coli* ECM properties on biofilm morphology and mechanical properties.

Independent of the scenario of their assembly, the resulting fibrous composite material is expected to swell due to water uptake by the pEtN-cellulose. While such swelling-induced expansion preferably occurs towards the surface interface of the biofilm, i.e. rather vertically than laterally, it may also be constrained by the tight interactions with the stiff curli fibers (**Figure 6b**). The fibrous matrix would then undergo stress-induced orientation in the direction of the swelling. Such mechanical explanation could justify the vertical orientation of the fibrous matrix observed in cross-sections of biofilms containing both pEtN-cellulose and curli (**Figure 2 and 6b**), also described as pillars in the literature,^2,3^ which appear more marked where interactions between pEtN-cellulose and curli are maximum (AR3110 and co-seeded 50:50).

This representation of the interaction between curli and pEtN-cellulose fibers in *E. coli* biofilms further aligns with findings obtained at the highest hierarchical level, i.e. at the biofilm scale (**Figure 6c**). Indeed, the similarities in mesoscopic architecture among the biofilms containing both fibers (**Figure 1a and 2**), modulated by the differences in their pEtN-cellulose content (and thus swelling capacity) (**Figure 1f**) is consistent with the coefficients characterizing higher wrinkling in biofilms rich in pEtN-cellulose (**Figure S62c**). Moreover, the higher rigidity and lower apparent plasticity of AR3110 and co-seeded 50:50 biofilms concord with the vertical ECM fiber orientation resulting from constrained swelling and with the structural role of water in the viscoelastic behavior of the resulting composite (**Figure 1b-c**).

## Conclusion

The growth of *E. coli* AR3110 biofilms involves the co-production and tight association of rigid curli and swelling pEtN-cellulose fibers: the two main building blocks of their ECM.^8,29^ This work showed that the co-assembly of these components with complementary properties dictates the structure and biophysical characteristics of the supramolecular product, which also appeared to be structurally independent of its components, i.e. a composite material. Spectroscopy experiments and electronic microscopy performed on fibers assembled by the same (AR3110) or different bacteria (co-seeded 50:50) indicated that slight differences in the conditions of assembly of the building blocks have a significant impact on the nanostructure and morphology of the matrix (composite vs hybrid material), but not so much at the biofilm scale. In addition, this study also suggests that the structural and mechanical properties of the resulting *E. coli* matrix composite vary depending on the ratio of building blocks pEtN-cellulose to curli, which can be tuned by co-seeding the corresponding *E. coli* strains in different ratio (AP329 and W3110 respectively).

Overall, this work suggests that it is possible to take advantage of the composite nature of bacterial matrix to reach a large range of combinations of mechanical and biophysical properties by directing both the proportion of the involved building blocks and their interactions. Such knowledge can benefit the development of ELMs with desired properties, whereby synthetic biology used to program the nature of the building blocks can be combined with various strategies to tune their ratio and assembly conditions. This approach could be extended to other bacterial biofilm producers but also to fungi and algae, and introduced as a new strategy to control the properties of bio-sourced materials.

## Experimental Section

### 1. Bacterial strain and growth

This work used different *E. coli* K-12 strains derived from *E. coli* K-12 W3110, a strain synthesizing amyloid curli protein but not pEtN-cellulose.^13^ **Table 2** describes each strain used:

**Table 2.** Description of the strains used in this study, summarized from Siri et al.^16^.

| Strain name | Extracellular matrix | Comment |
| --- | --- | --- |
| AR3110 | pEtN-cellulose+:curli+ | Replacement of stop codon in *bcsQ* by a sense codon |
| W3110 | pEtN-cellulose-:curli+ | Stop codon in *bcsQ* |
| AP329 | pEtN-cellulose+:curli- | AR3110 *csgBA::kan* |

The *E. coli* strain AR3110 is a derivative of W3110 with a restored capacity to produce phosphoethanolamine (pEtN)-modified cellulose in addition to producing curli amyloid fibers as majors ECM components.^13,15^ AP329 (csgBA::kan) is an AR3110 derivative that is deficient in the production of curli while producing pEtN-cellulose.^13,15^ The *E. coli* strains AP329 and W3110 were mixed in different proportions at the time of seeding based on their OD600nm. Considering a ratio AP329:W3110, the proportions used in the co-seeded biofilms were 75:25, 50:50, and 25:75.

Salt-free Luria−Bertani (LB) agar plates (15 mm diameter) were prepared with 1.8 % w/v of bacteriological grade agar−agar (Roth, 2266), supplemented with 1 % w/v tryptone (Roth, 8952) and 0.5 % w/v yeast extract (Roth, 2363). After agar pouring, the plates were left to dry for 10 minutes with the lid open and 10 minutes with the lid partially open to avoid future condensation. Each agar plate was left to rest at room temperature for 48 hours before bacteria seeding. A suspension of bacteria was prepared from a single colony and grown overnight in LB medium at 37 °C with shaking at 250 rpm. Each plate was inoculated with arrays of 9 drops of 5 μL of bacterial suspension (OD_600nm_ ∼ 0.5 after 10x dilution). After inoculation, the excess of water evaporated from the drops and left bacteria-rich dry traces. Biofilms were grown for 5 days (∼ 120 h) inside an incubator at 28 °C. The relative humidity in the incubator remained around 30 %RH.

### 2. Biofilm imaging

Six biofilms per condition were imaged with a stereomicroscope (AxioZoomV.16, Zeiss, Germany) using the tiling function of the acquisition software (Zen 2.6 Blue edition, Zeiss, Germany). To estimate the biofilm projected area, 6 independent biofilms were measured at their equatorial line using the Fiji software.^47^ An average was then calculated for each growth condition.

### 3. Biofilm growth kinetics

For the biofilm growth kinetics, a live imaging experiment of the biofilm growth was conducted.^11^ B biofilms were grown in a custom-made on-stage incubator installed on the motorized stage of an AxioZoomV.16 stereomicroscope (Zeiss, Germany). At 1-hour intervals 3 × 3 and 2 × 2 tile regions were automatically recorded in brightfield at 4–9 positions on a 15 cm Petri dish over the course of 121 hours. Temperature and relative humidity inside the on-stage incubator were controlled and set to 28 °C and >90 %, respectively.

Biofilm spreading area A(t) was analyzed automatically using custom-written MATLAB codes (Matlab 9.7.0 R2019b, MathWorks, Natick, MA). In a first step, thresholding was applied to the intensity images to segment biofilm from the background. As image contrast increased due to biofilm growth, thresholding was possible from t > 10 h. For each condition, Ai = 1 was defined at the time point, when all different samples grown in this condition could be detected, to have a common reference point for the calculation of the relative area increase A(t)/Ai. A growth curve based on these results was then plotted using GraphPad Prism v9.0 software.

### 4. Biofilm water content, dry mass and water uptake

The water content and water uptake of the biofilms were determined by scraping 7 biofilms per condition from the respective agar substrates after 5 days of growth (∼ 120 hours). Biofilms were placed in plastic weighing boats, and dried at 60 °C for 3 hours in an oven. Wet and dry masses (m_wet_, m_dry_) were determined before and after drying.^11^ The biofilms water content in each growth condition was estimated with **Eq. (1)**

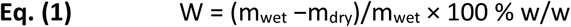

To determine the water uptake (W_up_), we added Millipure water in excess (5 mL) to the dehydrated biofilms from each condition, covered them with aluminum foils to avoid evaporation and left overnight at room temperature. The water excess was removed and the biofilm samples were weighed again (m_rewet_). The percentage of water uptake of biofilms after rehydration (%W_up,w_) was determined with respect to biofilm initial wet mass as described in **Eq. (2)**

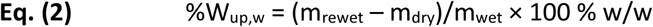

The water uptake per gram of dry biofilm (W_up,d_) was calculated with **Eq. (3)**

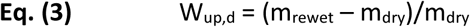

All procedures were carried out in four independent experiments.

### 5. Fluorescence confocal microscopy

For these experiments, biofilms were grown on salt-free agar supplemented with the fluorescent dye Direct Red 23 (Sigma-Aldrich, Germany, also known as Pontamine Fast Scarlett 4b) in a final concentration of 0.015 g/L. The protocol established to obtain cross sections of living biofilms was adapted from Ziege et al.^11^ Briefly, the biofilms of interest were isolated by trimming the underlying agar substrate slowly immersed with 50°C hot liquid salt-free agar without added nutrients. The resulting agar−biofilm−agar sandwiches were then cut into ∼ 1 mm thick slices using a blade, and placed in thin glass slides.

The slices of each biofilm were observed with a LEICA confocal microscope SP8 FALCON (Leica, Mannheim, Germany) with an oil immersion 63X objective (1.2NA) under excitation at 552 nm and collecting signal in the emission range 600-700 nm. Images were taken and analyzed with the LAS X software. Six cross-sections were imaged per biofilm.

Fiji software was also used to analyze the wrinkling coefficient.^47^ Each image was turned to an 8-bit image and the LUT was inverted, a threshold was established and then the length of the cross-section was measured using the straight-line tools, the measurement was defined as biofilm length (L_0_). Finally, the free-hand line tool was used to follow the path of wrinkles to define the wrinkles length (L_w_). The biofilm wrinkling ratio δ_w_ was calculated by **Eq. (4)**

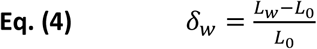

### 6. Micro-indentation on biofilms

1. *E. coli* biofilms of each strain were grown for 5 days on the same plate and used for the indentation experiments. A TI 950 Triboindenter (Hysitron Inc.) equipped with a conospherical tip (r=50 µm) was used to determine the load–displacement curves after calibration of the instrument in air. Ten measurements were performed in the central region of each biofilm. The distance between two measurement points was at least 250 μm in x and y directions and the depth of the indentation was between 10 and 25 µm, i.e. much less than biofilm the thickness (∼ 75 µm).^48^ Loading rates ranged from 20 μm/s, which translates to loading and unloading times of 10 s. The loading portion of all curves were fitted with a Hertzian contact model over and indentation range of 0 to 10 µm to obtain the reduced elastic modulus E_r_.^11,17^

An apparent plasticity index (ψ’) was defined as **Eq. (5)** (**Figure S3**)

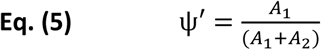

where A_1_ describes the area between the loading and unloading curves and A_2_ describes the area under the unloading curve. ψ’ spans between the values 0 – 1, where ψ’ = 1 characterizes a plastic behavior (irreversible deformation), and ψ’ = 0 indicates an elastic response (reversible deformation).

For the apparent holding plasticity (ψ’_h_), we added a holding step at maximum displacement of 10 s before the retracting steps. The ψ’_h_ was defined as **Eq. (6)** (**Figure S4**)

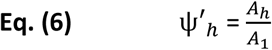

where A_h_ describes the area between the loading and unloading curves during the holding point, and A_1_ describes the area under the loading curve. The ψ’_h_ spans between 0 – 1, where ψ’_h_ = 0 means the biofilm is not able to dissipate energy, and ψ’_h_ = 1 means the biofilm can fully dissipate energy from the force applied by the indentation tip during the holding time.

A subset of curves with a maximum indentation depth of 20 µm were selected for the analysis of the adhesion force, and visco-elastic load relaxation.

The adhesion force *F*_*ad*_ is the minimum force measured during tip retraction.

For visco-elastic load relaxation, the corresponding load-time curves from 0 - 10 s were extracted with constant displacement. From each curve, we calculated the portion of the force relaxation happening before a tip instability as occurring at 205ms as described in Siri et al^15^ following **Eq. (7)**

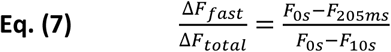

where Δ*F*_*fast*_ represents the fast relaxation reaction occurring between t= 0 – 205 ms, and define *F*_*total*_ as the total force relaxed after 10 s (endpoint for the holding time).

### 7. Curli fiber purification

Fiber purification involved a similar process as reported in previous works.^31^ Briefly, a total of 27 biofilms (∼ 1 g of biofilm material) were scraped from the surface of the substrates. Biofilms were blended five times on ice with an XENOX MHX 68500 homogenizer for 1 min at 2-min intervals. The bacteria were pelleted by centrifuging two times at low speed (5000g at 4°C for 10 min). A final concentration of NaCl 150 mM was added to the supernatant and the curli pelleted by centrifuging at 12.000g at 4°C for 10 min. The pellet was resuspended in 1mL of solution containing 10 mM tris (pH 7.4) and 150 mM NaCl, and incubated on ice for 30 min before being centrifuged at 16.000g at 4°C for 10 min. This washing procedure was repeated thrice. The pellet was then resuspended in 1mL of 10 mM tris solution (pH 7.4) and pelleted as described above (16.000 g at 4°C for 10 min). The pellet was again suspended in 1 mL of 10 mM tris (pH 7.4) and centrifuged at 17.000 g at 4°C for 10 min. This washing step was repeated twice. The pellet was then resuspended in 1 mL of SDS 1 % v/v solution and incubated for 30 min. The fibers were pelleted by centrifuging at 19.000 g at 4°C for 15 min. The pellet was resuspended in 1 mL of Milli-Q water. This washing procedure was repeated thrice. The last resuspension was done in 0.1 mL of Milli-Q water supplemented with 0.02 % sodium azide. The fiber suspension was stored at 4°C for later use.

### 8. Transmission electron microscopy (TEM)

2 μL drops of fiber suspension were adsorbed onto Formvar-coated carbon grids (200 mesh), washed with Milli-Q water, and stained with 1 % w/v uranyl acetate. The samples were imaged in a JEOL-ARM F200 transmission electron microscope equipped with two correctors for imaging and probing. For the observations, we applied an acceleration voltage of 200 kV. The width of the purified fibers values in TEM images were measured using the scale tool of the GATAN GMS 3 software. To avoid subjectivity over 10 different images and over 10 identified fibers per field were used.

### 9. Cryo-Focused Ion-Beam

The different ratios of purified fibers were pipetted between two type A gold-coated copper freezer hats (BALTIC preparation, Wetter, Germany) and rapidly frozen using the ICE high-pressure freezing system (Leica Microsystems, Vienna, Austria). For the different areas within the biofilms, the latter were initially fixed using paraformaldehyde (PFA 4 %) for 2 hours. Following fixation, a thin layer of agar was applied to encapsulate the biofilms, making it easier to isolate the regions of interest (ROIs). These ROIs were carefully cut using a scalpel and placed between two type B gold-coated copper freezer hats (BALTIC preparation, Wetter, Germany), with Hexadecene (Sigma-Aldrich) added to act as a cryo-protectant. The hats were arranged in a mirrored configuration to have a cavity thickness of 0.6 mm. Samples were maintained at liquid nitrogen temperatures and transferred to a cryo-holder in the Leica EM VCM loading station (Leica Microsystems, Vienna, Austria). Using the VCT500 shuttle, they were moved to the ACE600 system (Leica Microsystems, Vienna, Austria) for freeze-fracturing and metal coating. Fractured surfaces were coated first with a 10 nm carbon layer, then an 8 nm layer of platinum. The prepared specimens were subsequently transferred via the VCT500 shuttle to the Zeiss Crossbeam 540 (Zeiss Microscopy GmbH, Oberkochen, Germany) for imaging. Throughout all handling steps, sample temperatures were kept below −145°C. For all fibers and biofilms ROIs samples, a trench of approximately 30 μm length and 60 μm width was milled at 30 nA with the ion beam. The exposed surface was polished and imaged with a reduced beam current of 1.5 nA. Imaging via FIB-SEM Serial Surface View was conducted with the electron beam operating at 2 keV and a current of either 50 or 90 pA using a mixed of Inlens and secondary electron (SE) detectors. Images were acquired in a sequential “slice and view” manner, with the pixel resolution set to 8 nm in both x and y axes. The z-axis, or slice thickness, was also defined at 8 nm, ensuring uniform, isometric voxel dimensions throughout data acquisition.

### 10. Image processing and segmentation

Stacks alignment, image processing, and segmentations were done using Dragonfly software, Version 2024.1 (Object Research Systems (ORS) Inc, Montreal, Canada). All stacks were first automatically aligned using the sum of square differences (SSD) algorithm available in the slice registration module. To enhance structural clarity, image noise was minimized through the application of a convolution filter. Fibers segmentation, whether from purified samples or the ROIs within biofilms, was completed using the segmentation wizard in Dragonfly. For each dataset, eight slices were used to train the segmentation model, followed by manual refinement using the brush tool and the define threshold. In biofilms datasets, bacterial structures were also segmented using the same wizard, with further refinement and separation performed via the watershed algorithm.

### 11. Fluorescence spectroscopy

Corrected steady-state emission spectra were acquired with a FluoroMax®-4 spectrofluorometer (HORIBA). Spectra were recorded at 25°C using a 3-mm path cuvette (Hellma® Analytics). Thioflavin T (ThioT) (Sigma Aldrich, Germany) measurements were performed in samples containing 4 µL of each purified product, 1 mM ThioT in Glycine buffer, pH 8.2, using λ_exc_ = 446 nm and spectral bandwidths of 10 nm. Seven µL of each sample was diluted to a final volume of 45 µL for the intrinsic fluorescence spectra (Trp emission). These were acquired using λ_exc_= 280 nm and 5/5 nm slit bandwidths.

### 12. Circular Dichroism (CD) spectroscopy

Spectra of 10 µL of the purified components from biofilms in Milli-Q water were recorded with a Chirascan CD spectrometer (Applied Photophysics, Leatherhead, Surrey, UK) using a quartz cuvette with 1-mm path length (Hellma, Müllheim, Germany). Spectra were acquired between 190 nm and 250 nm wavelengths, with 1 nm step size, 1 nm band-width and 0.7 s integration time per point. Milli-Q water was used to define the measurement background, which was automatically subtracted during acquisition. The experiments were repeated thrice for each condition. Each measurement was an average of three scans.

### 13. Attenuated total reflectance Fourier transform infrared spectroscopy (ATR-FTIR)

IR spectra were acquired on a spectrophotometer (Vertex 70v, Bruker Optik GmbH, Germany) equipped with a single reflection diamond reflectance accessory continuously purged with dry air to reduce water vapor distortions in the spectra. Fibers in Milli-Q water samples (∼ 4 μL) were spread on a diamond crystal surface, dried under N2 flow to obtain the protein spectra. A total of 64 accumulations were recorded at 25 °C using a nominal resolution of 4 cm^−1^.

Spectra were processed using Kinetic software developed by Dr. Erik Goormaghtigh at the Structure and Function of Membrane Biology Laboratory, Université Libre de Bruxelles, Brussels, Belgium. After subtraction of water vapor and side chain contributions, the spectra were baseline corrected and area normalized between 1800 and 800 cm^−1^. For a better visualization of the overlapping components arising from the distinct structural elements, the spectra were deconvoluted using Lorentzian deconvolution factor with a full width at the half maximum (FWHM) of 20 cm^−1^ and a Gaussian apodization factor with a FWHM of 30 cm^−1^ to achieve a line narrowing factor K = 1.5.^49^ Second derivative was performed on the Fourier self-deconvoluted spectra for further analysis.

### 14. Protein quantification in purified fibers

Protein quantification of the purified fibers was done using a Bradford assay. Briefly, in a 96-well microplate, 5 µL of purified fibers from each biofilm was incubated with 195 µL of Bradford reagent (Quick Start™ Bradford 1x Dye Reagent #5000205, BioRad). Absorbance was measured at 550 nm after a 5–10 min incubation protected from the light. BSA (heat shock fraction, protease free, fatty acid free, essentially globulin free, pH 7, ≥ 98%, (A7030) Sigma) was used in different concentrations as a calibration curve. The experiments were repeated with three independent purification batches of each fiber.

### 15. Phasor analysis

Spectral phasor plots were used to visualize Nile Red bound to the fibers’ spectral shifts. The phasor plots are 2D scatter graphs, where the axes are the real (G) and imaginary (S) components of the Fourier transform of the fluorescence spectra. This transformation offers a powerful, model free, graphical method to characterize spectral^50^ and lifetime information.^51^ A detailed analysis can be found elsewhere.^45,52,53^ In this work, Nile Red spectra can be transformed using the following for x and y coordinates:

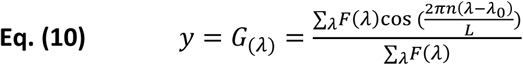

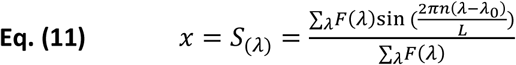

where F(λ) are the fluorescence intensity values, λ_0_ is the initial wavelength of the spectrum (λ_0_ =400 nm), L is the length of the spectrum and n is the harmonic value. The values L=300 (from 400 to 700 nm) and n=1 were used for phasor calculations. The angular position of the phasor in the plot is related to the center of mass, while the radial position depends on the full width at half maximum of the spectrum. Spectral phasor plots were constructed using Originlab data analysis software.

### 16. Fluorescence Lifetime Imaging Microscopy (FLIM)

Two-photon excitation of Nile red (NR) was performed at 860 nm with a pulsed Ti:Sapphire laser with 80Hz repetition rate (Spectra-Physics Mai Tai, Mountain View, CA).^17^ The image size was 512x512 pixels, the pixel size was 75 nm and the detection range were 600 to 728 nm. FLIM calibration of the system was performed by measuring the known lifetime of the fluorophore Coumarin 6 in ethanol.^53^ For these experiments, 5 μL fibers were placed in clean glass and 5 μM of NR was added.

The FLIM phasor analysis allows the transformation of the fluorescence signal from each pixel in the image to a point in the phasor plot. FLIM data were processed using SimFCS, an open source software developed at the Laboratory of Fluorescence Dynamics, Irvine, California (available at http://www.lfd.uci.edu).

The fitting of FLIM decay curves was performed with the integrated FLIM analysis software of the Leica SP8 microscope using the n-exponential reconvolution method. Three lifetimes were used to fit the curves obtaining chi-squared values close to one. The mean values weighted by intensity are informed.

### 17. Cryo–ESEM biofilm microscopy

Two biofilms (AR3110 and 50:50 co-seeded) were fixed in PFA and a layer of agar was applied on top to aid in handling and preserve structural integrity. For freezing, sections of ≈2x2x10 mm were cut from the two agar-coated biofilms and one side was gently immersed into a liquid nitrogen bath. This approach allowed the top layer to freeze first, promoting directional solidification and enhancing overall cooling efficiency while minimizing thermal shock. Once fully frozen, the samples were mounted into a cryo-holder under liquid nitrogen in the Leica EM VCM loading station (Leica Microsystems GmbH, Wetzlar, Germany). Subsequently, the samples were transferred using the VCT500 shuttle to the ACE600 system (Leica Microsystems, Vienna, Austria), where they were freeze fractured and sputter coated with a 10 nm layer of platinum. Finally, the coated samples were shuttled to the ESEM Quattro S (Thermo Scientific, Eindhoven, The Netherlands) for cryo-imaging. Images were acquired under cryogenic conditions using the secondary electron detector at an electron energy of 5 keV, with a working distance ranging from 5 to 6 mm. Samples were kept below -142 °C for the entire experiment once frozen.

### 18. Statistical analysis

For each experiment, 3 to 4 fiber solutions were used, where each solution came from different fiber purification batches. For each purification, 27 biofilms for each strain/mixture were cultured, and the different samples of fibers obtained from each purification were treated simultaneously (or in consecutive days) to minimize unavoidable variability in the implementation of the protocols (e.g. temperature and humidity in the laboratory during agar preparation and/or biofilm seeding).

For statistical analysis, a Shapiro Wilk test was used to check for data normality. For data with no normal distribution, Kruskal-Wallis non-parametric test was performed. For data with normal distribution, a One-way ANOVA test was carried out. Mechanical properties data was analyzed using a Mann-Whitney U test. Details of each test are described in the legend of the figures.

## Supporting Information

The following files are available free of charge:

- Biofilm morphology: Figure S1, Figure S2
- Biofilm mechanics: Figure S3, Figure S4, Figure S5 and Table S1 – 5
- Biofilm cross-section analysis: Figure S6 and Table S6
- ATR-FTIR spectroscopy: Fiber second derivative: Figure S7 and Table S7
- Protein concentration estimation in purified fibers: Figure S8
- Average lifetimes and standard deviations: Table S8

## Data Availability Statement

The data that support the findings of this study are available from the corresponding authors upon reasonable request.

## Author Contributions

Conceptualization: C.M.B and M.S. Visualization: C.M.B and M.S. Methodology: C.M.B, M.S, and E.R. Investigation: M.S, A.M, A.S, S.A, N.R and E.R. Writing Original Draft: M.S and C.M.B. Review and Editing: all authors. Supervision: C.M.B. Project Administration: C.M.B. Resources: C.M.B.

## Acknowledgements

M.S. acknowledges support from the Max Planck Queensland Centre on the Materials Science for Extracellular Matrices, and CONICET. A.M. acknowledges support from the AvH foundation and CONICET. C.M.B. thanks the financial support of the German Research Foundation (Deutsche Forschungsgemeinschaft ─ DFG) – within the research unit InterDent (FOR2804). The authors also thank Christine Pilz-Allen for her technical support in the laboratories and Heike Runge for her help in doing the transmission electronic microscopy experiments. The authors thank Regine Hengge and Diego Serra for the insightful discussions on the *E. coli* biofilm topic throughout the years. The authors are also grateful to Eric Goormaghtigh from the SFMB group at the Université Libre de Bruxelles for providing the Kinetics Software. Finally, we thank Kerstin Blank for helping in the transformation of the *E. coli* W3110 bacteria with the plasmid pMP7604 (gene for the fluorescent protein mCherry) kindly donated by Guido V. Bloemberg (University of Zurich).

## References

1. Wang, S., Zhan, Y., Jiang, X. & Lai, Y. Engineering Microbial Consortia as Living Materials: Advances and Prospectives. ACS Synth Biol 10.1021/acssynbio.4c00313 (2024) doi:10.1021/acssynbio.4c00313.

2. An, B. et al. Engineered Living Materials For Sustainability. Chem Rev 123, 2349–2419 (2023).

3. Nguyen, P. Q., Courchesne, N. M. D., Duraj-Thatte, A., Praveschotinunt, P. & Joshi, N. S. Engineered Living Materials: Prospects and Challenges for Using Biological Systems to Direct the Assembly of Smart Materials. Advanced Materials 30, 1–34 (2018).

4. Wang, Y., Zhang, Q., Ge, C., An, B. & Zhong, C. Programmable Bacterial Biofilms as Engineered Living Materials. Acc Mater Res 5, 797–808 (2024).

5. Mekontso, J. A., Farrukh, U., Trujillo, S. & del Campo, A. A practical workflow for cytocompatibility assessment of living therapeutic materials. Biomaterials Advances 169, (2025).

6. Wilking, J. N., Angelini, T. E., Seminara, A., Brenner, M. P. & Weitz, D. A. Biofilms as complex fluids. MRS Bull 36, 385–391 (2011).

7. Serra, D. O. & Hengge, R. A c-di-GMP-Based Switch Controls Local Heterogeneity of Extracellular Matrix Synthesis which Is Crucial for Integrity and Morphogenesis of Escherichia coli Macrocolony Biofilms. J Mol Biol 431, 4775–4793 (2019).

8. Thongsomboon, W. et al. Phosphoethanolamine cellulose: A naturally produced chemically modified cellulose. Science (1979) 359, 334–338 (2018).

9. Serra, D. O. & Hengge, R. Stress responses go three dimensional - The spatial order of physiological differentiation in bacterial macrocolony biofilms. Environ Microbiol 16, 1455–1471 (2014).

10. Yan, J., Nadell, C. D., Stone, H. A., Wingreen, N. S. & Bassler, B. L. Extracellular-matrix-mediated osmotic pressure drives Vibrio cholerae biofilm expansion and cheater exclusion. Nat Commun 8, (2017).

11. Ziege, R. et al. Adaptation of Escherichia coli Biofilm Growth, Morphology, and Mechanical Properties to Substrate Water Content. ACS Biomater Sci Eng 7, 5315–5325 (2021).

12. McCrate, O. A., Zhou, X., Reichhardt, C. & Cegelski, L. Sum of the Parts: Composition and Architecture of the Bacterial Extracellular Matrix. J Mol Biol 425, 4286–4294 (2013).

13. Serra, D. O., Richter, A. M. & Hengge, R. Cellulose as an architectural element in spatially structured escherichia coli biofilms. J Bacteriol 195, 5540–5554 (2013).

14. O., S. D., M., R. A. & Regine, H. Cellulose as an Architectural Element in Spatially Structured Escherichia coli Biofilms. J Bacteriol 195, 5540–5554 (2013).

15. Siri, M. et al. Mechanical Comparison of Escherichia coli Biofilms with Altered Matrix Composition: A Study Combining Shear-Rheology and Microindentation. ACS Biomater Sci Eng 11, 4523–4536 (2025).

16. Jeffries, J. et al. Variation in the ratio of curli and phosphoethanolamine cellulose associated with biofilm architecture and properties. Biopolymers 112, 1–11 (2021).

17. Siri, M., Mangiarotti, A., Vázquez-Dávila, M. & Bidan, C. M. Curli fibers in Escherichia coli biofilms: the influence of water availability on amyloid structure and properties. Macromol Biosci 2022.11.21.517345 (2023) doi:doi.org/10.1002/mabi.202300234.

18. Siri, M., Vazquez-Davila, M., Guzman, C. S. & Bidan, C. M. Nutrient availability in fl uences E. coli bio fi lm properties and the structure of puri fi ed curli amyloid fi bers. NPJ Biofilms Microbiomes 10, (2024).

19. Flemming, H. C. & Wingender, J. The biofilm matrix. Nat Rev Microbiol 8, 623–633 (2010).

20. Tyrikos-Ergas, T. et al. Synthetic phosphoethanolamine-modified oligosaccharides reveal the importance of glycan length and substitution in biofilm-inspired assemblies. Nat Commun 13, 3954 (2022).

21. Horvat, M., Pannuri, A., Romeo, T., Dogsa, I. & Stopar, D. Viscoelastic response of Escherichia coli biofilms to genetically altered expression of extracellular matrix components. Soft Matter 15, 5042–5051 (2019).

22. Cegelski, L. et al. Small-molecule inhibitors target Escherichia coli amyloid biogenesis and biofilm formation. Nat Chem Biol 5, 913–919 (2009).

23. Serra, D. O. & Hengge, R. Cellulose in Bacterial Biofilms BT - Extracellular Sugar-Based Biopolymers Matrices. in (eds. Cohen, E. & Merzendorfer, H.) 355–392 (Springer International Publishing, Cham, 2019). doi:10.1007/978-3-030-12919-4_8.

24. Axpe, E. et al. Fabrication of Amyloid Curli Fibers-Alginate Nanocomposite Hydrogels with Enhanced Stiffness. ACS Biomater Sci Eng 4, 2100–2105 (2018).

25. Khatri, V. et al. Bionanocomposites with Enhanced Physical Properties from Curli Amyloid Assemblies and Cellulose Nanofibrils. Biomacromolecules 24, 5290–5302 (2023).

26. Serra, Di O. & Hengge, R. Bacterial Multicellularity: The Biology of Escherichia coli Building Large-Scale Biofilm Communities. Annu Rev Microbiol 75, 269–290 (2021).

27. Wang, Q. & Zhao, X. A three-dimensional phase diagram of growth-induced surface instabilities. Sci Rep 5, 1–10 (2015).

28. Klauck, G., Serra, D. O., Possling, A. & Hengge, R. Spatial organization of different sigma factor activities and c-di-GMP signalling within the three-dimensional landscape of a bacterial biofilm. Open Biol 8, (2018).

29. Eva, B., Andrea, B., Corinne, D. & Paolo, L. Gene Expression Regulation by the Curli Activator CsgD Protein: Modulation of Cellulose Biosynthesis and Control of Negative Determinants for Microbial Adhesion. J Bacteriol 188, 2027–2037 (2006).

30. Siri, M., Herrera, M., Moyano, A. J. & Celej, M. S. Influence of the macromolecular crowder alginate in the fibrillar organization of the functional amyloid FapC from Pseudomonas aeruginosa. Arch Biochem Biophys 713, 109062 (2021).

31. Chapman, M. R. et al. Role of Escherichia coli curli operons in directing amyloid fiber formation. Science (1979) 295, 851–855 (2002).

32. Brown, E. E. & Laborie, M. P. G. Bioengineering bacterial cellulose/poly(ethylene oxide) nanocomposites. Biomacromolecules 8, 3074–3081 (2007).

33. Dayal, M. S. & Catchmark, J. M. Mechanical and structural property analysis of bacterial cellulose composites. Carbohydr Polym 144, 447–453 (2016).

34. Arkharova, N. A. et al. SEM and TEM for structure and properties characterization of bacterial cellulose/hydroxyapatite composites. Scanning 38, 757–765 (2016).

35. Sleutel, M., Pradhan, B., Volkov, A. N. & Remaut, H. Structural analysis and architectural principles of the bacterial amyloid curli. Nat Commun 14, 1–14 (2023).

36. Biancalana, M. & Koide, S. Molecular mechanism of Thioflavin-T binding to amyloid fibrils. Biochim Biophys Acta Proteins Proteom 1804, 1405–1412 (2010).

37. Mishra, R. et al. Natural Anti-biofilm Agents: Strategies to Control Biofilm-Forming Pathogens. Front Microbiol 11, (2020).

38. Krebs, M. R. H., Bromley, E. H. C. & Donald, A. M. The binding of thioflavin-T to amyloid fibrils: Localisation and implications. J Struct Biol 149, 30–37 (2005).

39. Mishra, R., Sjölander, D. & Hammarström, P. Spectroscopic characterization of diverse amyloid fibrils in vitro by the fluorescent dye Nile red. Mol Biosyst 7, 1232–1240 (2011).

40. Barth, A. Infrared spectroscopy of proteins. Biochim Biophys Acta Bioenerg 1767, 1073–1101 (2007).

41. Bertasa, M. et al. A study of non-bounded/bounded water and water mobility in different agar gels. Microchemical Journal 139, 306–314 (2018).

42. Lakowicz, J. R. Principles of Fluorescence Spectroscopy, 3rd Principles of Fluorescence Spectroscopy, *Springer, New York, USA*, 3rd *Edn*, 2006. Principles of fluorescence spectroscopy*, Springer, New York, USA, 3rd edn*, *2006.* (2006). doi:10.1007/978-0-387-46312-4.

43. Levitt, J. A., Chung, P.-H. & Suhling, K. Spectrally resolved fluorescence lifetime imaging of Nile red for measurements of intracellular polarity. J Biomed Opt 20, 096002 (2015).

44. Golini, C. M., Williams, B. W. & Foresman, J. B. Further Solvatochromic, Thermochromic, and Theoretical Studies on Nile Red. J Fluoresc 8, 395–404 (1998).

45. Malacrida, L., Ranjit, S., Jameson, D. M. & Gratton, E. The Phasor Plot: A Universal Circle to Advance Fluorescence Lifetime Analysis and Interpretation. Annu Rev Biophys 50, 575–593 (2021).

46. Torrado, B., Malacrida, L. & Ranjit, S. Linear Combination Properties of the Phasor Space in Fluorescence Imaging. Sensors vol. 22 Preprint at 10.3390/s22030999 (2022).

47. Schindelin, J., et al. Fiji: An open-source platform for biological-image analysis. Nat Methods 9, 676–682 (2012).

48. Serra, D. O., Klauck, G. & Hengge, R. Vertical stratification of matrix production is essential for physical integrity and architecture of macrocolony biofilms of Escherichia coli. Environ Microbiol 17, 5073–5088 (2015).

49. Goormaghtigh, E., Cabiaux, V. & Ruysschaert, J.-M. Secondary structure and dosage of soluble and membrane proteins by attenuated total reflection Fourier-transform infrared spectroscopy on hydrated films. Eur J Biochem 193, 409–420 (1990).

50. Fereidouni, F., Bader, A. N. & Gerritsen, H. C. Spectral phasor analysis allows rapid and reliable unmixing of fluorescence microscopy spectral images. Opt Express 20, 12729 (2012).

51. Jameson, D. M., Gratton, E. & Hall, R. D. The measurement and analysis of heterogeneous emissions by multifrequency phase and modulation fluorometry. Appl Spectrosc Rev 20, 55–106 (1984).

52. Ranjit, S., Malacrida, L., Jameson, D. M. & Gratton, E. Fit-free analysis of fluorescence lifetime imaging data using the phasor approach. Nat Protoc 13, 1979–2004 (2018).

53. Digman, M. A., Caiolfa, V. R., Zamai, M. & Gratton, E. The phasor approach to fluorescence lifetime imaging analysis. Biophys J 94, 14–16 (2008).

