## Supplementary Information for "*E. coli* extracellular matrix: a tunable composite with hierarchical structure"

### Supporting Information

| | |
|---|---|
| Biofilm morphology | Figure S1 |
| | Figure S2 |
| Biofilm mechanics | Figure S3 |
| | Figure S4 |
| | Figure S5 |
| | Table S1 – S5 |
| Biofilm cross-section analysis | Figure S6 |
| | Table S6 |
| ATR-FTIR spectroscopy: Fiber second derivative | Figure S7 |
| | Table S7 |
| Protein concentration estimation in purified fibers | Figure S8 |
| Average lifetimes and standard deviations | Table S8 |

### 1. *Biofilm morphology*

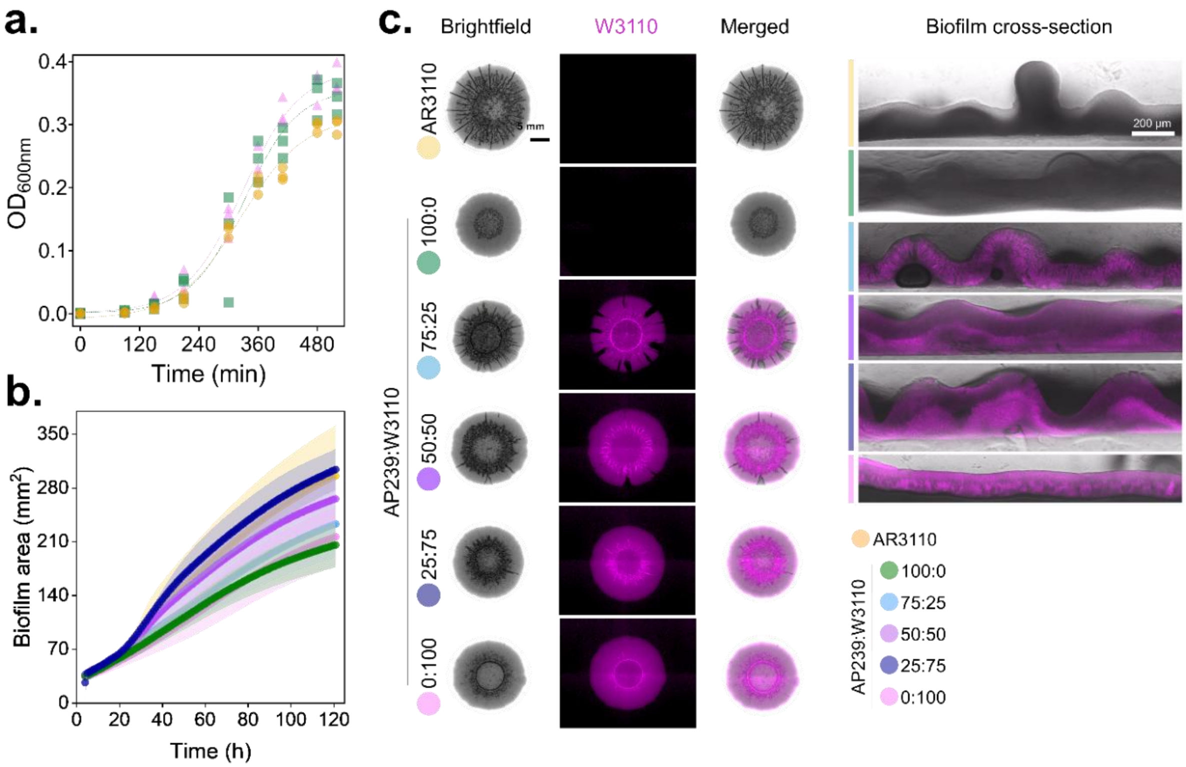


**Figure S 1 *E. coli* biofilm morphology and bacterial arrangement of different strains.** N=6 (a) Bacterial growth curve of the different *E. coli* strains. (b) Biofilm growth kinetics of the mixed and pure biofilms over 5 days (121 h). To see representative videos of the growth kinetics of each biofilm please see the supplementary videos. (c) Stereomicroscopies of the *E. coli* biofilms in this study. W3110 bacteria (curli producing bacteria) are m-cherry labelled to see the bacteria distribution in the biofilms (left column). Confocal biofilm cross-sections of the imaged biofilms in the stereomicroscope (right column). Independent experiments were done in 3-4 biofilms, except otherwise stated.

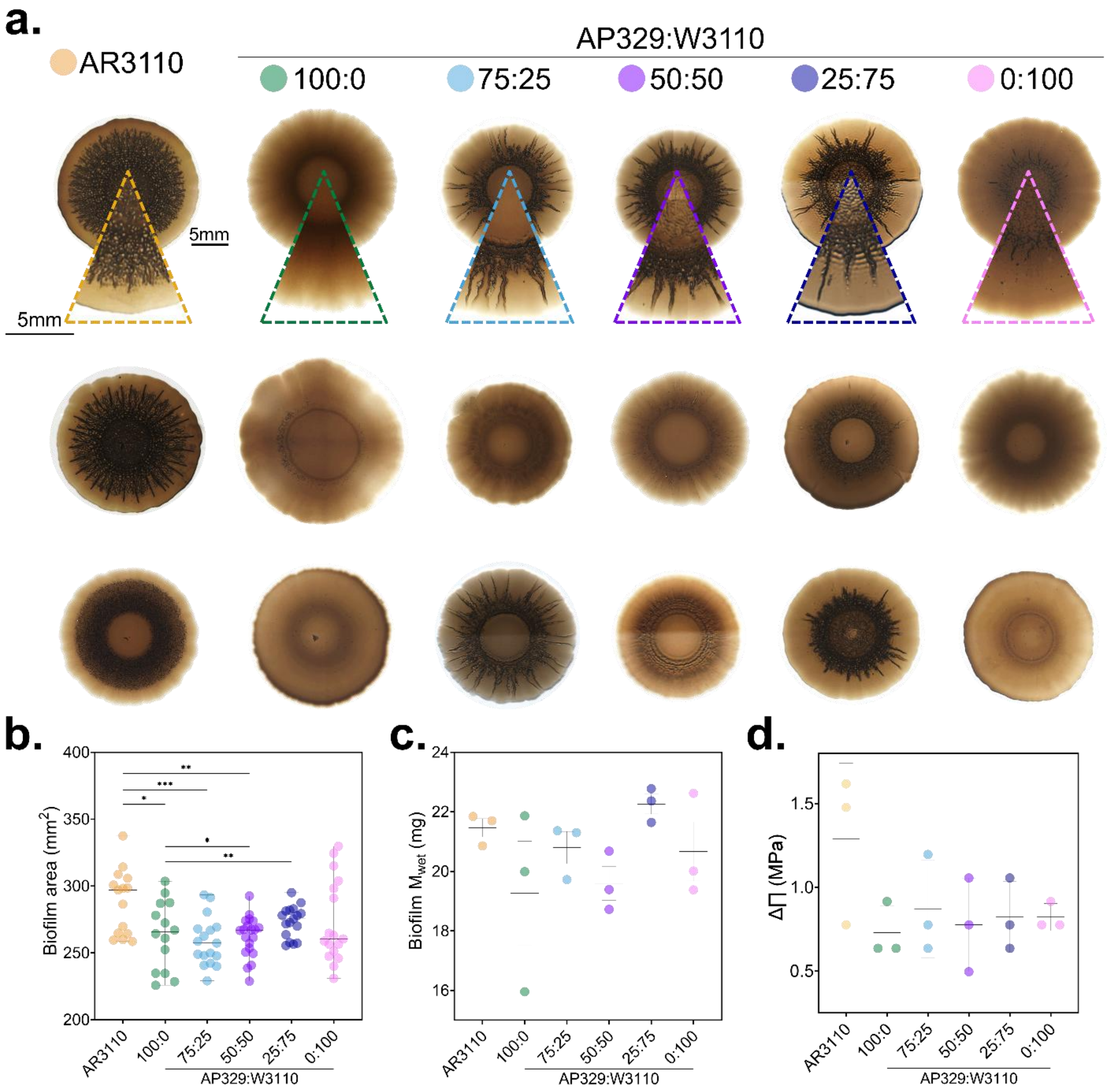


**Figure S 2 General characteristics of *E. coli* biofilms from different strains**. (a) Stereomicroscopies of the variability of the morphology of *E. coli* biofilms of different strains. Scale bar = 5 mm. (b) Biofilm area. The statistical analysis was done with Man Whitney U test (p<0.0001, **** | p<0.001, *** | p<0.01, ** | p<0.05, * | ns = non-significant). Data acquired from 15 - 20 independent biofilms per condition. (c) Biofilm wet mass after scraping them from the surface. (d) Osmotic gradient between the biofilm and the agar substrate. All data presented here come from N=4 independent biofilm cultures for each condition tested, and the statistical analysis was done with One-way ANOVA (p<0.001, *** | p<0.01, ** | p<0.05, * | ns = non-significant).

### 2. *Biofilm mechanics*

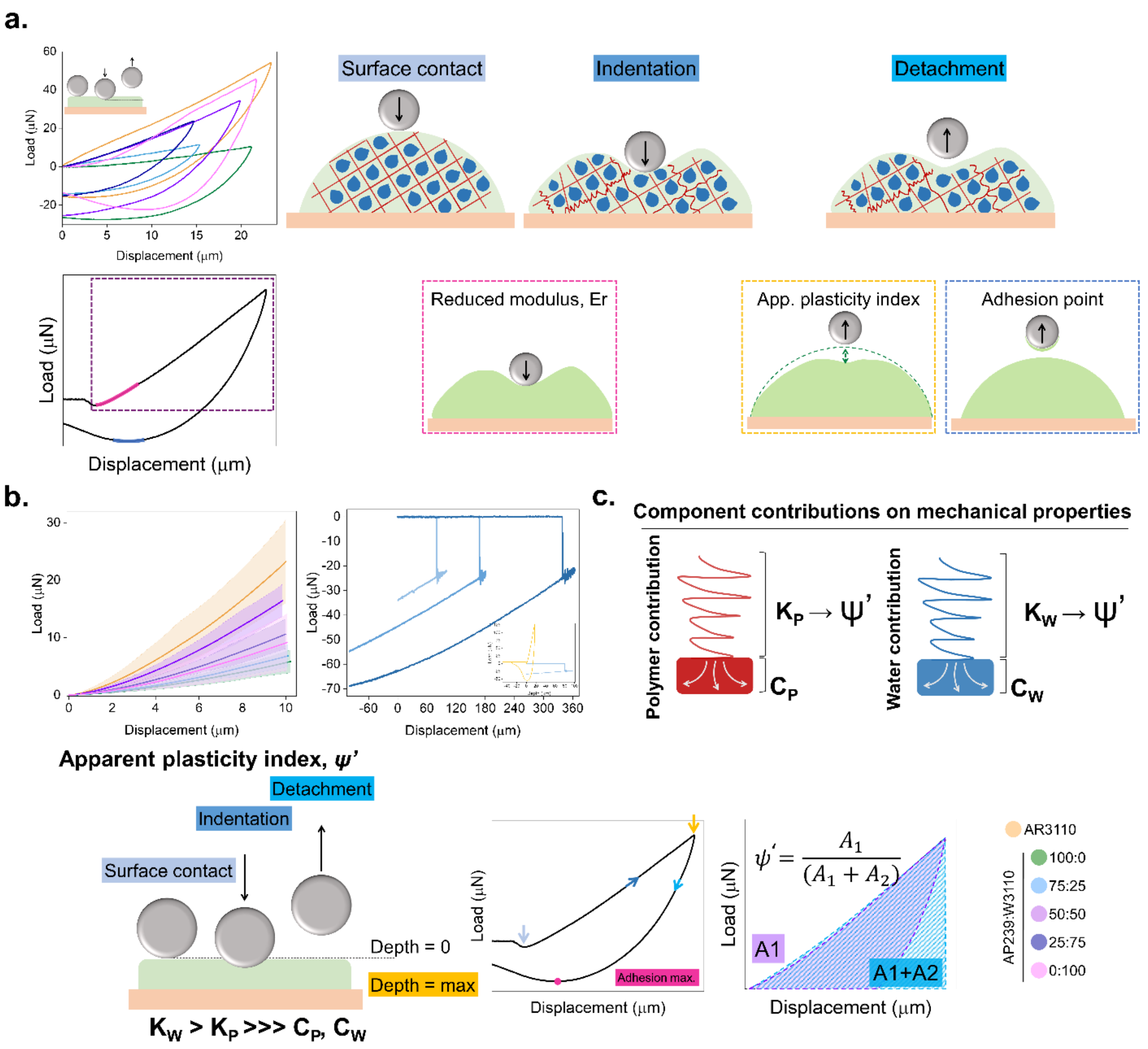


**Figure S 3 Microindentation "short-time scale" experiment design and control for the *E. coli* biofilms.** (a) Representative load displacement curves when indenting the biofilm surface with a spherical tip indentation with a diameter of 50 µm, and scheme representation of the experiment set-up. (b) Zoom in on part of the loading curves used for fitting a Hertzian contact model. Range of the fitting curves for the reduced modulus estimation of each biofilm studied in this study. Considering the size of the tip (conospherical tip, r=50 µm) and thickness of the biofilms (100 µm), the displacement range taken into account for the reduced modulus was ~10 µm (first panel). Control indentation curves of water at different displacement maxima revealing that the tip attraction and adhesion forces in tip-water contacts are significantly larger than those between the tip and the biofilms. An inset was included comparing an AR3110 indentation curve, against a water indentation curve (second panel). (c) Contributions of the extracellular components of the biofilms in the apparent plasticity indexes with schematic details of the estimation of the different indexes calculated in the different loading-displacement curves.

The main components of the extracellular matrix that contribute to their mechanical properties are namely, polymers (curli amyloid fibers in our *E. coli*), and water. When performing micro-indentation experiments, upon contact with the biofilm, the film generates a force that may deform irreversibly the area where contact was made.

The response to this force varies depending on the reaction of the polymers and water to it. We can describe in a simple way the reaction of both components as a spring attached to its base (**Figure S3-4a**). The spring portion of the response informs about the apparent plasticity index (K) of said element; how elastic its behavior is in presence of the applied force. The base portion of the component's response informs about the dissipation index (C); how easily can the component relax from the force applied.

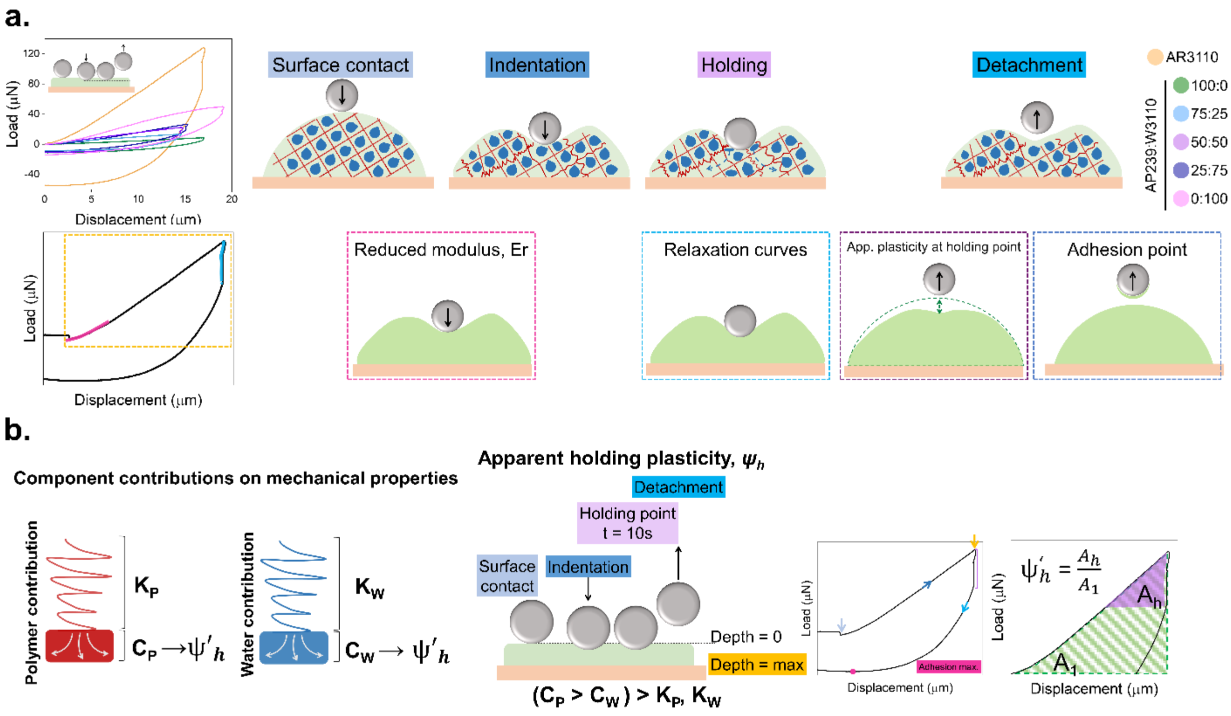


**Figure S 4 Microindentation "long-time scale" experiment design and control for the *E. coli* biofilms.** (a) Representative load displacement curves when indenting the biofilm surface with a spherical tip indentation with a diameter of 50 µm, and scheme representation of the experiment set-up. (b) Contributions of the extracellular components of the biofilms in the apparent plasticity at holding point with schematic details of the estimation of the different indexes calculated in the different loading-displacement curves.

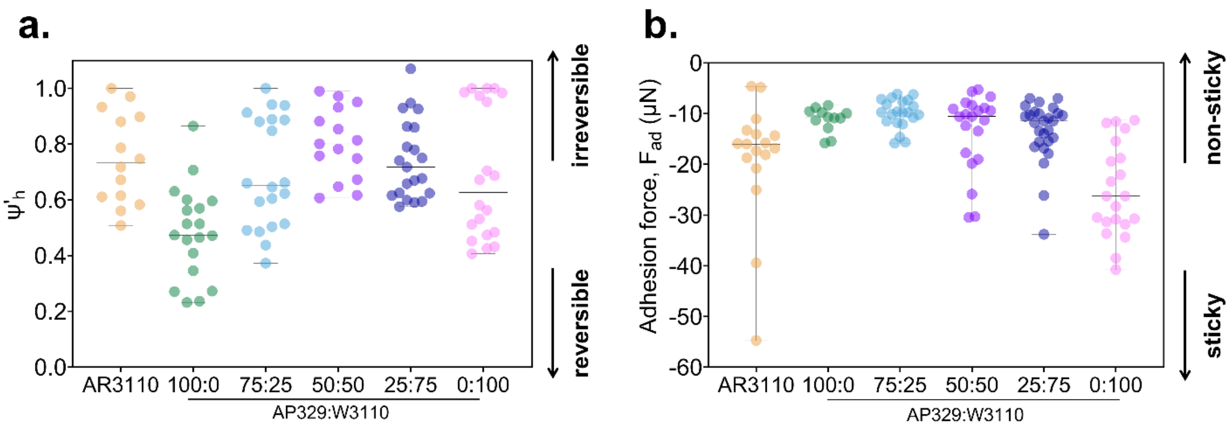

**Figure S 5 Mechanical characterization of the different strains of *E. coli* biofilms with microindentation.** (a) Apparent plasticity index at holding point, $\psi'_h$ defined as capacity of the biofilm to dissipate the energy from the force applied by the indentation tip after the end of the holding point (Figure S3). (b) Adhesion force, $F_{ad}$ measured from the minimum load recorded during tip retraction. Data obtained correspond to indentation curves with a maximum depth of 20 µm. N = 3 different biofilms. The statistical analysis was done with Man Whitney U test (p<0.00=1, **** | p<0.001, *** | p<0.01, ** | p<0.05, * | ns = non-significant) (Table S4- S5).

We studied both portion of behavior by carrying out two different experimental set-ups. For the apparent plasticity index, we indented the tip until the maximum depth and then started its retraction – detachment step (**Figure S3a**). This set-up allows us to measure the first response of the material to the force applied by the indenting tip (the elastic properties of the material). For the apparent holding plasticity, we indented the tip until the maximum depth, and held the tip in that position for 10 seconds before starting the detachment step (**Figure S4a**). During the 10 seconds in which the tip was in the material, the different components of the biofilm matrix had time to relax and adjust to this new force applied. The apparent holding plasticity reports on how much the material could reorganize around the tip. These indexes where then calculated as described in the experimental section.

**Table S 1** The statistical analysis was done with Man Whitney U test (p<0.00=1, **** | p<0.001, *** | p<0.01, ** | p<0.05, * | ns = non-significant) from the data in Figure 4a.

**Biofilm stiffness (kPa)**

| STRAINS | AR3110 | 100:0 | 75:25 | 50:50 | 25:75 | 0:100 |
|---|---|---|---|---|---|---|
| AR3110 | | **** | **** | **** | **** | **** |
| 100:0 | **** | | ** | **** | **** | * |
| 75:25 | **** | ** | | **** | **** | ns |
| 50:50 | **** | **** | **** | | **** | **** |
| 25:75 | **** | **** | **** | **** | | *** |
| 0:100 | **** | * | ns | **** | *** | |

**Table S 2** The statistical analysis was done with Man Whitney U test (p<0.00=1, **** | p<0.001, *** | p<0.01, ** | p<0.05, * | ns = non-significant) from the data in Figure 4b.

**Apparent plasticity index, ψ'**

| STRAINS | AR3110 | 100:0 | 75:25 | 50:50 | 25:75 | 0:100 |
|---|---|---|---|---|---|---|
| AR3110 | | **** | **** | ns | ns | ** |
| 100:0 | **** | | ** | **** | **** | *** |
| 75:25 | **** | ** | | *** | **** | **** |
| 50:50 | ns | **** | *** | | ns | ** |
| 25:75 | ns | **** | **** | ns | | **** |
| 0:100 | ** | *** | **** | ** | **** | |

**Table S 3** The statistical analysis was done with Man Whitney U test (p<0.00=1, **** | p<0.001, *** | p<0.01, ** | p<0.05, * | ns = non-significant) from the data in Figure 4c.

**$\Delta F_{fast}/\Delta F_{total}$**

| STRAINS | AR3110 | 100:0 | 75:25 | 50:50 | 25:75 | 0:100 |
|---|---|---|---|---|---|---|
| AR3110 | | **** | ns | ns | ns | Ns |
| 100:0 | **** | | ns | **** | *** | ns |
| 75:25 | ns | ns | | * | ns | ns |
| 50:50 | ns | **** | * | | ns | * |
| 25:75 | ns | *** | ns | ns | | ns |
| 0:100 | ns | ns | ns | * | ns | |

**Table S 4** The statistical analysis was done with Man Whitney U test (p<0.00=1, **** | p<0.001, *** | p<0.01, ** | p<0.05, * | ns = non-significant) from the data in Figure 4e.

**Apparent plasticity at holding point, $\psi'_h$**

| STRAINS | AR3110 | 100:0 | 75:25 | 50:50 | 25:75 | 0:100 |
|---|---|---|---|---|---|---|
| AR3110 | | ns | ns | ns | ns | ns |
| 100:0 | ns | | ns | ** | ns | ns |
| 75:25 | ns | ns | | ns | ns | ns |
| 50:50 | ns | ** | ns | | ns | ns |
| 25:75 | ns | ns | ns | ns | | ns |
| 0:100 | ns | ns | ns | ns | ns | |

**Table S 5** The statistical analysis was done with Man Whitney U test (p<0.00=1, **** | p<0.001, *** | p<0.01, ** | p<0.05, * | ns = non-significant) from the data in Figure 4f.

**Adhesion force, $F_{ad}$ (μN)**

| STRAINS | AR3110 | 100:0 | 75:25 | 50:50 | 25:75 | 0:100 |
|---|---|---|---|---|---|---|
| AR3110 | | ** | *** | ns | * | * |
| 100:0 | ** | | ns | ns | ns | **** |
| 75:25 | *** | ns | | ns | * | **** |
| 50:50 | ns | ns | ns | | ns | **** |
| 25:75 | * | ns | * | ns | | **** |
| 0:100 | * | **** | **** | **** | **** | |

### 3. *Biofilm cross-section analysis*

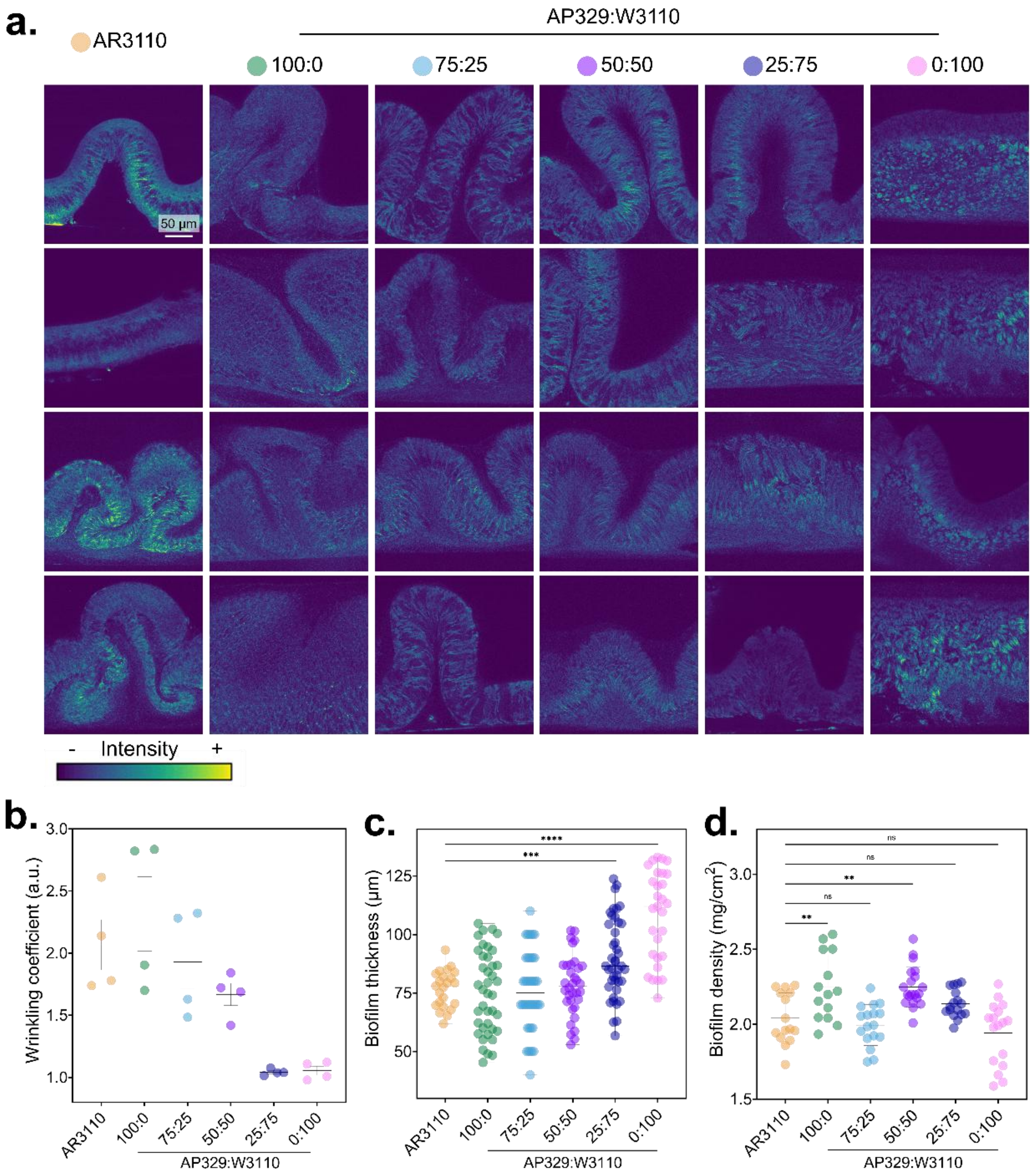


**Figure S 6 Biofilm cross-section under confocal imaging.** (a) Biofilm zoomed crossections from Figure 2 depicting the Direct Red intensity. (b) Wrinkling quantification. The wrinkling coefficient for each biofilm was calculated from confocal cross-sections of the different biofilms like the representative images in Figure 2. See materials and methods for details on the estimation. (c) Biofilm matrix thickness. (d) Estimation of biofilm density based on the biofilm dry mass.

**Table S 6** One-way AONVA statistical analysis of Figure S3; (p<0.001, *** | p<0.01, ** | p<0.05, * | ns = non-significant) with a Tukey's post-test for multiple comparisons (alpha=0.05) comparing all samples against all samples. N=6

**Wrinkling coefficient $\delta_w$**

| STRAINS | AR3110 | 100:0 | 75:25 | 50:50 | 25:75 | 0:100 |
|---|---|---|---|---|---|---|
| AR3110 | | ns | ns | ns | ** | ** |
| 100:0 | ns | | ns | ns | *** | *** |
| 75:25 | ns | ns | | ns | * | * |
| 50:50 | ns | ns | ns | | ns | ns |
| 25:75 | ** | *** | * | ns | | ns |
| 0:100 | ** | *** | * | ns | ns | |

*4. ATR-FTIR spectroscopy: Fiber second derivative*

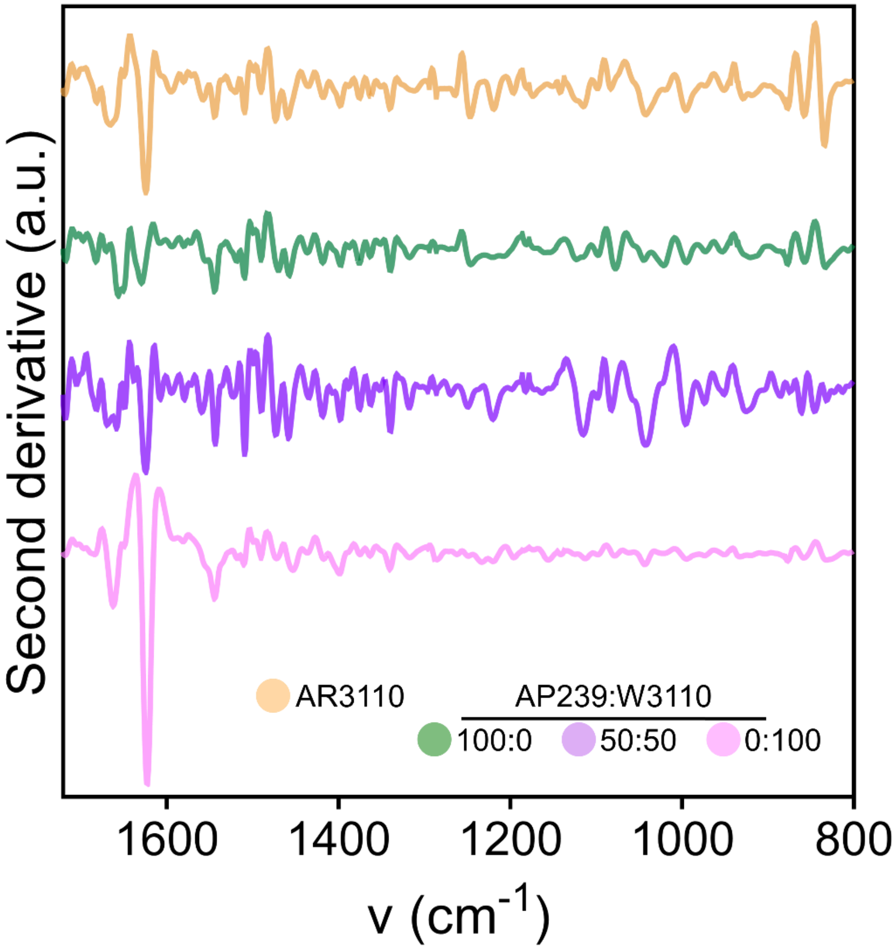


**Figure S 7 Second derivative of the ATR-FTIR spectra of the purified fibers in Figure 6c**

**Table S 7** One-way AONVA statistical analysis of Figure 6; (p<0.001, *** | p<0.01, ** | p<0.05, * | ns = non-significant) with a Tukey's post-test for multiple comparisons (alpha=0.05) comparing all samples against all samples. N=4

**$Abs_{curli}/Abs_{pEtN\text{-}cellulose}$**

| STRAINS | AR3110 | 100:0 | 75:25 | 50:50 | 25:75 | 0:100 |
|---|---|---|---|---|---|---|
| AR3110 | | ns | ns | ns | ns | *** |
| 100:0 | ns | | ns | ns | ns | ** |
| 75:25 | ns | ns | | ns | ns | *** |
| 50:50 | ns | ns | ns | | ns | *** |
| 25:75 | ns | ns | ns | ns | | * |
| 0:100 | *** | ** | *** | *** | * | |

### *5. Protein concentration estimation in purified fibers*

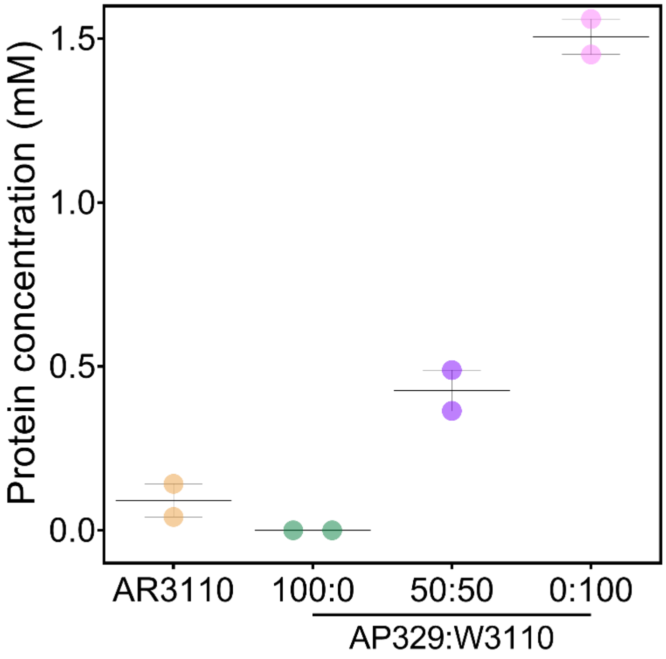


**Figure S 8 Protein concentration in the purified fibers.** Estimation of the curli component concentration in the purified fibers. The estimation was done by Bradford's test of the purified fibers after purification as described in Siri et al. [2] The statistical analysis was done with One-way ANOVA (p<0.0001, **** | p<0.001, *** | p<0.01, ** | p<0.05, * | ns = non-significant), post test used was Tukey's test to compare each fiber against every fiber.

### *6. Average lifetimes and standard deviations*

**Table S 8** Average lifetimes components used for fitting the curves in Figure 5h.

| Strains | | Lifetime 1 | SD | Lifetime 2 | SD | Lifetime 3 | SD |
|---|---|---|---|---|---|---|---|
| *AR3110* | | **0,46** | 0,09 | **1,52** | 0,49 | **3,98** | 0,29 |
| *AP329:W3110* | *100:0* | **0,27** | 0,16 | **0,96** | 0,20 | **3,19** | 0,07 |
| | *50:50* | **0,41** | 0,02 | **1,35** | 0,09 | **2,85** | 0,06 |
| | *0:100* | **0,41** | 0,02 | **1,25** | 0,08 | **3,05** | 0,10 |

### References


1. Salem, K. S. *et al.* Comparison and assessment of methods for cellulose crystallinity determination. *Chem. Soc. Rev.* **52**, 6417–6446 (2023).

2. Siri, M., Vazquez-Davila, M., Guzman, C. S. & Bidan, C. M. Nutrient availability in fl uences E . coli bio fi lm properties and the structure of puri fi ed curli amyloid fi bers. *npj Biofilms Microbiomes* **10**, (2024).

3. Wang, Q. & Zhao, X. A three-dimensional phase diagram of growth-induced surface instabilities. *Sci. Rep.* **5**, 1–10 (2015).
